# Intricate response dynamics enhances stimulus discrimination in the resource-limited *C. elegans* chemosensory system

**DOI:** 10.1101/2024.01.14.575365

**Authors:** Eduard Bokman, Christian O. Pritz, Rotem Ruach, Eyal Itskovits, Hadar Sharvit, Alon Zaslaver

## Abstract

Sensory systems evolved intricate designs to accurately encode perplexing environments. However, this encoding task may become particularly challenging for animals harboring a small number of sensory neurons. Here, we studied how the compact resource-limited chemosensory system of *C. elegans* uniquely encodes a range of chemical stimuli. We find that each stimulus is encoded using a small and unique subset of neurons, where only a portion of the encoding neurons sense the stimulus directly, and the rest are recruited via inter-neuronal communication. Furthermore, while most neurons show stereotypical response dynamics, some neurons exhibit versatile dynamics that are either stimulus specific or network-activity dependent. Notably, it is the collective dynamics of all responding neurons which provides valuable information that ultimately enhances stimulus identification, particularly when required to discriminate between closely-related stimuli. Together, these findings demonstrate how a compact and resource-limited chemosensory system can efficiently encode and discriminate a diverse range of chemical stimuli.

## Introduction

Living organisms critically rely on chemical signals. These signals direct fundamental behaviors such as locating food sources and mating partners, or avoiding toxins and predators. Sensory systems therefore evolved to differentiate between the multitude of chemical stimuli to allow animals to form an accurate representation of the environment (Prasad and Reed 1999; Zufall and Munger 2016).

In vertebrates, chemosensation is segregated into the olfactory and the gustatory modalities. Each of these modalities is relaying the information through several neural layers, and this distributed information is then integrated in deeper cortical areas (Small and Green 2012). Within the olfactory system itself, accurate identification of odorants is performed already in the glomeruli and relies on both the population coding and the timing in which each olfactory neuron was activated (Wilson et al. 2017; Chong et al. 2020).

Invertebrates, particularly those with small nervous systems, have shallower networks with limited neural layers. As such, chemosensory information is likely to be integrated, even partially, already at the level of the sensory layer (Bargmann 2006; Sengupta 2007; Ferkey, Sengupta, and L’Etoile 2021; Ghosh et al. 2017; Yu, Xue, and Chen 2022; Metaxakis, Petratou, and Tavernarakis 2018). But how can such sensory systems with limited sensory resources uniquely encode many different stimuli?

For example, the nervous system of *C. elegans* nematodes consists of 302 neurons with well-established synaptic connections (White et al. 1986; Cook et al. 2019; Witvliet et al. 2021). The main chemosensory organ, the amphid, is situated anteriorly and includes 12 bilaterally symmetrical pairs of neurons. Many of these chemosensory neurons respond to a variety of chemical stimuli, including olfactory and gustatory cues (Zaslaver et al. 2015; Lin et al. 2023; Yemini et al. 2021; Suzuki et al. 2008; How, Navlakha, and Chalasani 2021). Indeed, single cell RNA-seq data indicate that each neuron expresses an array of chemosensory receptors (Hammarlund et al. 2018; Taylor et al. 2021).

The *C. elegans* connectome shows a high degree of inter-connectivity within the sensory layer, suggesting that sensory coding may be shaped by neural communication among the sensory neurons or via feedback from the downstream interneurons (Chalasani et al. 2010). While some of the neurons respond to the stimulus directly (primary neurons), others respond through recruitment via chemical synapses, electrical gap junction, or humorally via neuropeptide signaling. This unique representation poses even greater coding limitations since only a smaller subset, consisting of the primary neurons only, can uniquely encode the target stimulus.

Ample studies quantified neural responses to a myriad of stimuli, focusing mainly on the sensory population coding and the response magnitudes (Schrödel et al. 2013; Zaslaver et al. 2015; Yemini et al. 2021; How, Navlakha, and Chalasani 2021; Lin et al. 2023). These studies revealed that the sensory system employs a hierarchical sparse coding scheme, whereby some neurons respond to a wide range of stimuli, while others are more selective. In addition to population coding, response dynamics was also shown to be carrying important functional information. For example, pulsatile activity in the sensory neuron AWA underlies an efficient chemotactic navigation that allows animals to reach attractive cues faster (Itskovits et al. 2018; Ruach et al. 2022).

In this study, we comprehensively analyzed with cellular resolution how the chemosensory system of *C. elegans* responds to and codes various chemical stimuli. We reveal that stimuli are encoded using a small, usually bilaterally symmetric, subset of neurons, where primary neurons consist of only 2-4 neurons on average. Interestingly, some neurons possess rich response dynamics that is stimulus specific or network-communication dependent. This rich response dynamics repertoire significantly improves discernment between similar neural population coding, thus endowing limited sensory systems the capacity to encode a larger amount of information.

## Results

### A comprehensive functional analysis of the chemosensory system

To systematically study how the compact chemosensory system of *C. elegans* worms encodes various chemical stimuli, we imaged activity from virtually all of the chemosensory neurons. For this, we used a transgenic strain expressing the genetically-encoded calcium indicator GCaMP in all amphid sensory neurons (**Figure 1A-B**). Individual animals were inserted into a custom-made microfluidic device (Chronis, Zimmer, and Bargmann 2007), and neuronal activity was measured in response to diverse olfactory and gustatory stimuli, representing both attractive and repulsive agents: Isoamyl-alcohol (IAA), Diacetyl (DA), Sodium chloride (NaCl), hyperosmotic (1M) glycerol, Quinine, and Sodium dodecyl sulfate (SDS), where [IAA, DA, NaCl] are attractive cues, while [Glycerol, Quinine, SDS] are repellents. For all conditions, we assayed neural activity for both the presentation (ON step) and the removal (OFF step) of the stimulus (**Figure 1C**). To verify that the delivery, or the removal, of the stimulus was temporally accurate, we added a fluorescent dye (rhodamine) to the stimulus. We therefore also assayed neural responses to the buffer supplemented with rhodamine only (Control group, **Figure 1C**). In general, rhodamine alone elicited minimal responses and all our statistical analyses account for these background-level responses (See Methods).

**Figure 1.**
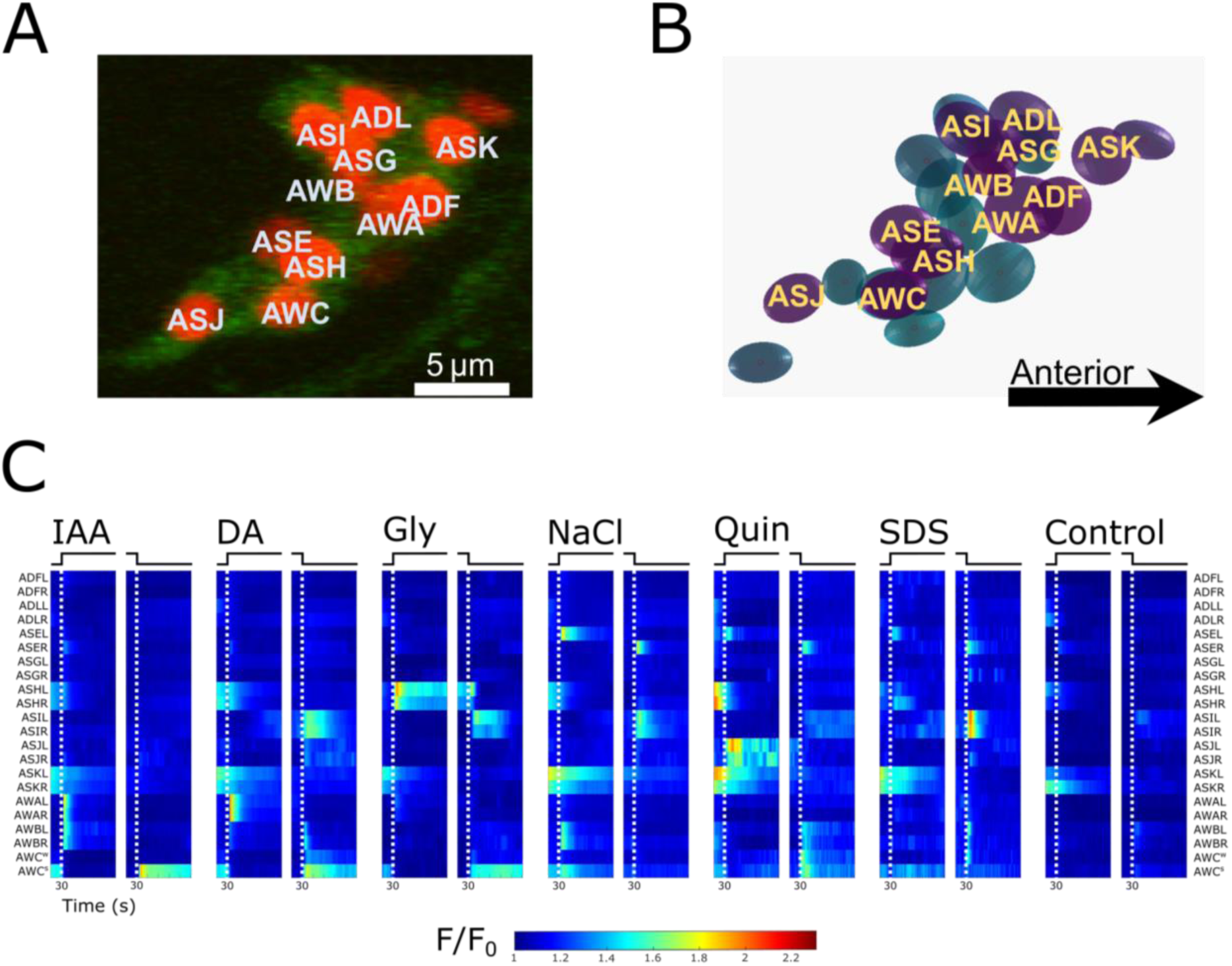
Functional dynamics of the *C. elegans* chemosensory system in response to a variety of chemical stimuli. **(A)** A confocal image of the right side of the amphid organ. Imaging was done using the strain *azrIs280* [*osm-6*::GCaMP3, *osm-6*::mCherry-NLS] (Iwanir et al. 2019; Pritz et al. 2023). Red, nuclear mCherry; Green, cytoplasmic GCaMP. Neuron identification relies on known anatomical position. **(B)** Visualization of the amphid nuclei segmented from (A). Fluorescence intensity was measured from the segmented spheres. Right side, purple; left side, blue. **(C)** Mean neural dynamics of individual neurons following stimulus presentation and removal. White dashed lines indicate ON/OFF steps. Note that the ASK, ASH and ADL neurons respond to blue light, hence the activity at the start of the imaging period. Conditions tested: Control (n=7); DA, diacetyl 10^-4^(n=23); IAA, isoamyl alcohol 10^-4^ (n=18); NaCl, sodium chloride 50mM (n=26); Glycerol 1M (n=7); Quinine 5mM (n=11); SDS 0.1% (n=12). A fluorescent red dye (500 nM rhodamine) was added to the stimuli to verify accurate stimulus switch. The control condition consisted of switching between buffer and buffer+dye. Responses observed in the control condition served as the baseline responses for neurons that may have responded to the dye only. The AWC pair is sorted by activation strength in each worm and is marked AWC^s^ (strong) and AWC^w^ (weak).

To simultaneously image all of the neurons, we used a confocal system equipped with a fast-resonating scanner that allowed imaging the entire brain volume at 2 Hz (30-40 slices, at a 0.6-0.7 µm Z-resolution), providing the necessary spatio-temporal resolution to reliably extract activity from individual sensory neurons (Schrödel et al. 2013; Nguyen et al. 2016; Randi and Leifer 2020; Iwanir et al. 2019; Yemini et al. 2021; Pritz et al. 2023; Lin et al. 2023). These acquisition settings, coupled with our analysis pipeline (see Methods) allowed tracking and measuring activity of all chemosensory neurons from both the right and left lateral sides (22 in total), excluding only AFDL/R which were often below detection levels (**Figure 1A-B**).

The populations of responding neurons per each stimulus were generally in line with previous reports (**Figure 1C**). For example, the AWC-type neurons responded to the removal of most stimuli in an OFF-step response manner (Chalasani et al. 2007). Similarly, ASH, known as polymodal aversive neurons (Kaplan and Horvitz 1993), responded upon encountering noxious stimuli, such as the hyperosmotic solution of 1M glycerol, and SDS. The ASEL and ASER neurons responded to the addition and removal of NaCl, respectively (Suzuki et al. 2008). Consistent with previous reports (Zaslaver et al. 2015; Lin et al. 2023), we observed a functional hierarchy in the chemosensory network where a few neurons appear to be general responders (AWC, AWB), responding to most or all of the tested stimuli, while other neurons are more selective to specific stimuli (e.g., ASG, AWA). All of the examined neuron classes responded to at least one of the tested conditions. Together, these findings indicate that individual neurons in this transgenic reporter strain are functionally intact, and that our automated analysis system reliably segments and identifies individual target neurons to extract accurate dynamic responses to various stimuli (**Figure 1C**).

### Lateral symmetric neurons generally show highly correlated activity

Apart from the ASE (Suzuki et al. 2008) and the AWC (Wes and Bargmann 2001) neurons, it is generally assumed that the left and right bilaterally symmetric sensory neurons exhibit similar neural responses (Lin et al. 2023; Zaslaver et al. 2015). We therefore utilized our ability to simultaneously measure functional responses from both lateral organs and analyzed the correlation between them. For this, we performed a pairwise correlation analysis between all the responding neurons across all six conditions.

Indeed, the lateral right- and left-symmetric neurons showed highly-correlated activity dynamics across all conditions, and the only exceptions to this were the AWC neurons, which responded asymmetrically in some conditions, and the ASER/L neurons that showed negative, or no correlation at all (**Figure 2)**. The correlation matrices also show that neuron types cluster in a stimulus-specific manner, as clustering varied across the different conditions **(Figure 2A)**. For example, in response to IAA, the activities of the AWA, AWB, and ASER neurons are correlated with each other, and negatively correlated with the activity of the AWC neurons. However, in response to Glycerol, activity of AWB and AWC is highly correlated, and negatively correlated with the activity of ASER. These results indicate a unique correlation pattern for each condition, providing a “finger-print” of the neuronal representation of a given stimulus. Due to the symmetry in responses, all neuron pairs, aside from AWC and ASE, were grouped for subsequent analyses.

**Figure 2.**
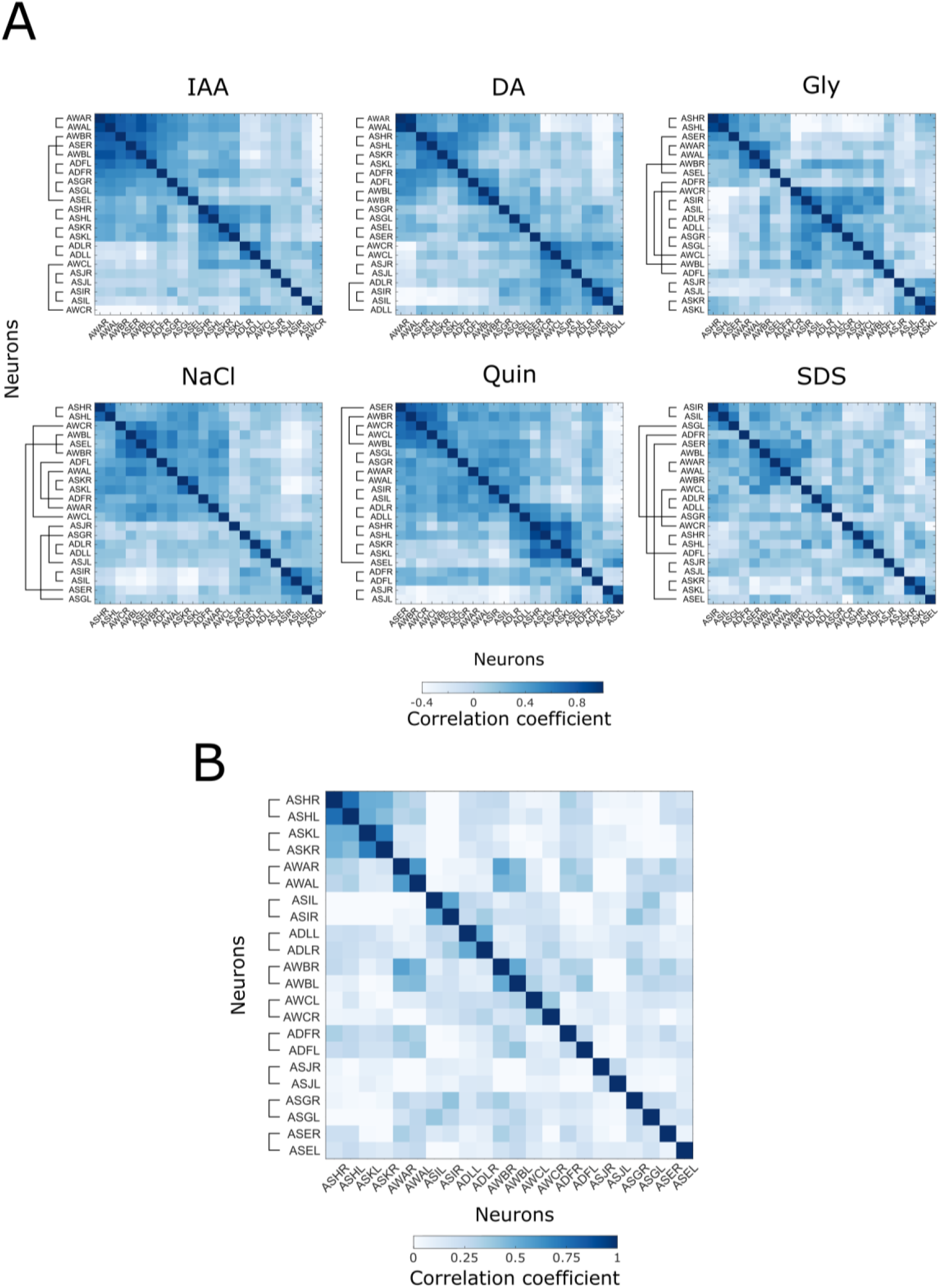
Activity of the right and left lateral symmetry neurons is highly correlated. **(A)** Pairwise time-series correlation matrices of the amphid neurons response dynamics. Correlations were first calculated across all neurons of each worm and then averaged over all worms in a condition. Pairs of right and left symmetric neurons are indicated by connecting lines. **(B)** Clustered activity across all conditions reveals the overall lateral functional symmetry. Only the ASER/L pair does not show correlated activity across all conditions.

### Neural dynamics varies in a stimulus-dependent manner

The strong correlation between the two lateral amphid neurons effectively reduces the number of “coding units” in the system by roughly a half. We therefore asked whether, in addition to the ensemble of responding neurons, stimulus identity could be further signaled by the activation dynamics of specific neurons.

Upon exposure to (or removal of) a stimulus, responding neurons typically show a sharp calcium increase that slowly, over several seconds, decays to baseline levels (**Figure 3**, blue). But how stereotypic are these response dynamics? For example, do individual neurons show stereotypic responses regardless of the specific stimulus? Do certain stimuli elicit the same response dynamics in different neurons? To address these questions, we performed a PC analysis on the response traces of all responding neurons across all of the conditions (**Figure 3**). The first two principal components combined explain ∼80% of the variance, and appear to reflect the absolute activity levels before and after presentation of the stimulus (**Supplementary figure s1A-B**). However, clustering by the PCs 3-4 (accounting for ∼10% of the variance) provides a clear separation that is based on the shape of the response dynamics. Most responses (∼75%) form a single cluster representing the stereotypical response dynamics of a sharp rise in Calcium levels to a narrow peak followed by an exponential-like decrease until resuming baseline levels (**Figure 3, blue**). This cluster includes each of the responding neurons in at least one condition, both ON and OFF step responses, and all tested stimuli.

**Figure 3.**
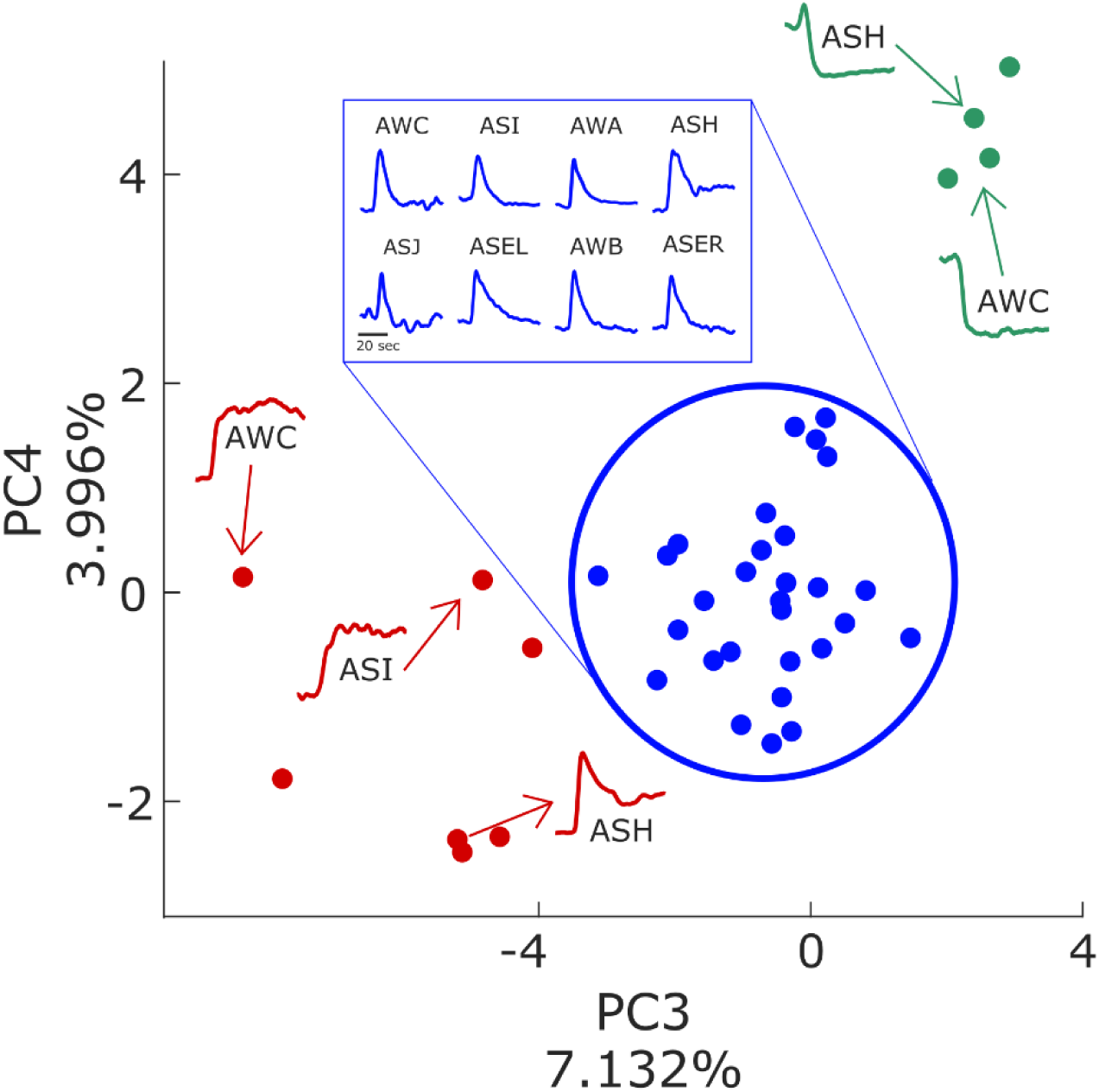
Activity dynamics varies in a stimulus-dependent manner. PC analysis of neuronal response dynamics. The PCA was performed on individual neuron traces, and each point is the average trace of a single neuron across all worms in a condition projected onto the PC space. K-means clustering revealed the dynamics differences in PCs three and four. The blue cluster represents stereotypical dynamics and includes examples from all neuron classes. The red and green clusters show non-stereotypical dynamics, and consist mostly of the ASH, ASI and AWC neurons in response to specific stimuli. Notably, these three neurons are also represented in the stereotypical-dynamics blue cluster. Inset traces show representative examples of response dynamics for each cluster. Each trace is normalized to its maximal level.

Two additional clusters represent variable response dynamics, including sustained elevated activity (AWC in IAA/DA OFF step. **Figure 3, red**), and inhibitory responses with decreased Calcium levels with (or without) an initial peak (ASH in Gly OFF step and AWC in IAA ON step, respectively, **Figure 3, green**). These variable responses were observed primarily in three neuron classes (AWC, ASH and ASI), suggesting that some neurons possess a larger repertoire of response dynamics than others, possibly providing more nuance in signaling stimulus identity.

Thus, while chemosensory neurons typically respond with very stereotypic activation dynamics, under some conditions, the same neurons exhibit vastly different dynamics. Such alternative responses suggest that sensory neurons may convey different messages depending on the specific stimulus, effectively increasing the information capacity of the sensory layer.

### Inter-neuronal communication shapes the sensory response

Sensory responding neurons can be classified as either primary responders, neurons that sense the stimulus directly (*e.g.* via a dedicated receptor), or secondary responders, neurons that are recruited to respond by other neurons (Leinwand and Chalasani 2013; Leinwand et al. 2015). This recruitment may be facilitated by synaptic neurotransmitters, extra-synaptically via secreted neuromodulators/neuropeptides, or through electrical gap junctions. The chemosensory neurons receive all these input types both laterally from sensory-layer neurons and from other neurons, most of which are interneurons (White et al. 1986; Varshney et al. 2011; Cook et al. 2019; Witvliet et al. 2021).

As internal communication may influence the response dynamics of individual neurons, we set out to discern the degree to which this inter-neuronal signaling shapes such sensory responses. For this, we measured response dynamics in *unc-13* and *unc-31* mutant strains that are defective in neurotransmitter (synaptic) and neuropeptide (extrasynaptic) release, respectively (**Figure 4A and supplementary figure 2**).

**Figure 4.**
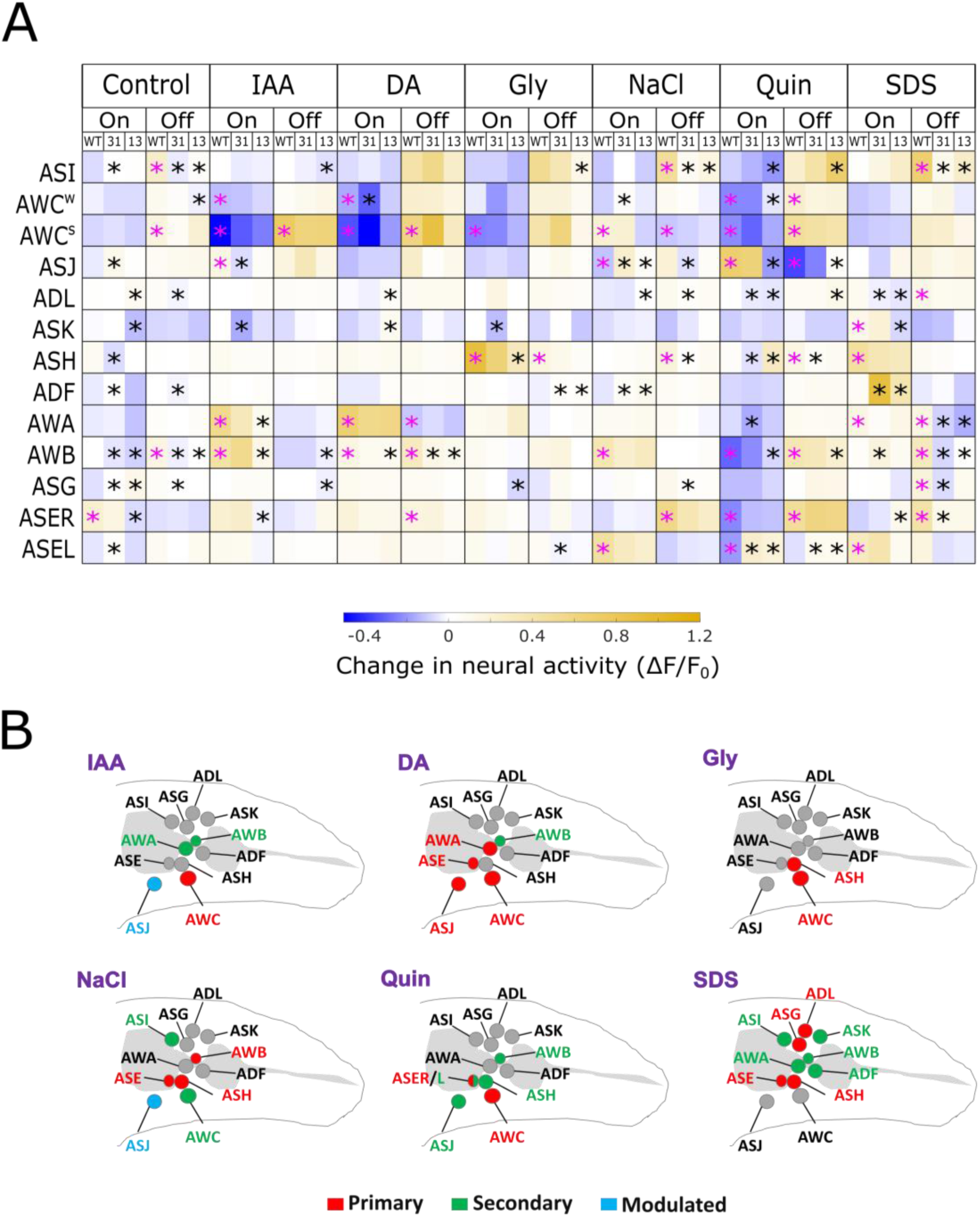
The *C. elegans* chemosensory system relies on extensive inter-neuronal communication. **(A)** Neural activities in response to ON and OFF steps of the different stimuli. Significant WT responses to each stimulus were compared to the control condition (pink asterisks, p<0.05), and significant mutant responses were compared to the WT response for the same condition (black asterisks, p<0.05). The neural activities are based on the first 7 seconds after each step (see Methods) Both sides of each neuron class were pooled, aside from the AWC and the ASE neurons. P-values were obtained using two-tailed t-tests corrected for multiple comparisons using FDR. **(B)** Schematic representation of Primary (red) and Secondary (green) responding neurons for each of the tested stimuli as determined by the dependence of the response on synaptic transmission. The ASJ neuron type, whose response was merely modulated (**Supplementary figure 3**), is denoted in cyan.

Comparing neural responses in these mutant strains to responses in WT animals reveals extensive inter-neuronal communication (**Figure 4A**). Over 40% of the neural responses showed altered activity in at least one of the mutants where they were either completely abolished or significantly diminished (marked in asterisks in figure 4A, and see concrete examples in **supplementary figure 3**). For example, the AWA and AWB neurons responded to IAA presentation in WT worms (pink asterisk), but failed to respond in the *unc-13* mutants (black asterisks), suggesting that these neurons are secondary responders to IAA and that they are recruited to respond via neurotransmitter signaling.

Subtler changes were observed in other neurons (e.g. the ASJ and ASH neurons) whose responses were merely modulated rather than completely abolished (**Supplementary figure s3**). Moreover, modulated activity of some neurons (e.g., ASJ) is stimulus specific and also depends on neuropeptide signaling (**Supplementary figure s3**). Dependence of neural activity on internal network signaling and on the specific stimulus may further increase the coding capacity of the sensory layer neurons.

Overall, out of the subset of responding neurons in each condition, only a few (typically 2-4) were unaffected by inter-neuronal communication and can therefore be classified as primary sensory neurons (**Figure 4B**). The secondary responders, forming the rest of the encoding ensemble, are recruited by the primary responders via inter-neuronal signaling, presumably to form the unique nuanced response of the specific stimulus.

### Stimulus identity can be predicted by network activity

If stimulus identity were only signaled by primary responders, the sensory system could face a combinatorial problem in that the variety of distinct environmental stimuli far outnumber the possible combinations of primary responders. To estimate how well population coding discriminates between the various stimuli, we trained a random forest classifier on the peak activities of the 13 chemosensory neurons (9 left-right pairs and the individual AWC and ASE neurons) in response to stimulus presentation and removal, for a total of 26 parameters per observation. The classifier perfectly predicted the identity of the stimulus presented to WT worms, and this prediction accuracy decreased the more neurons were removed from the training set (**Figure 5A and supplementary figure 4**). This suggests that when considering the entire ensemble of chemosensory neurons, it is very easy to discriminate between the different stimuli.

**Figure 5.**
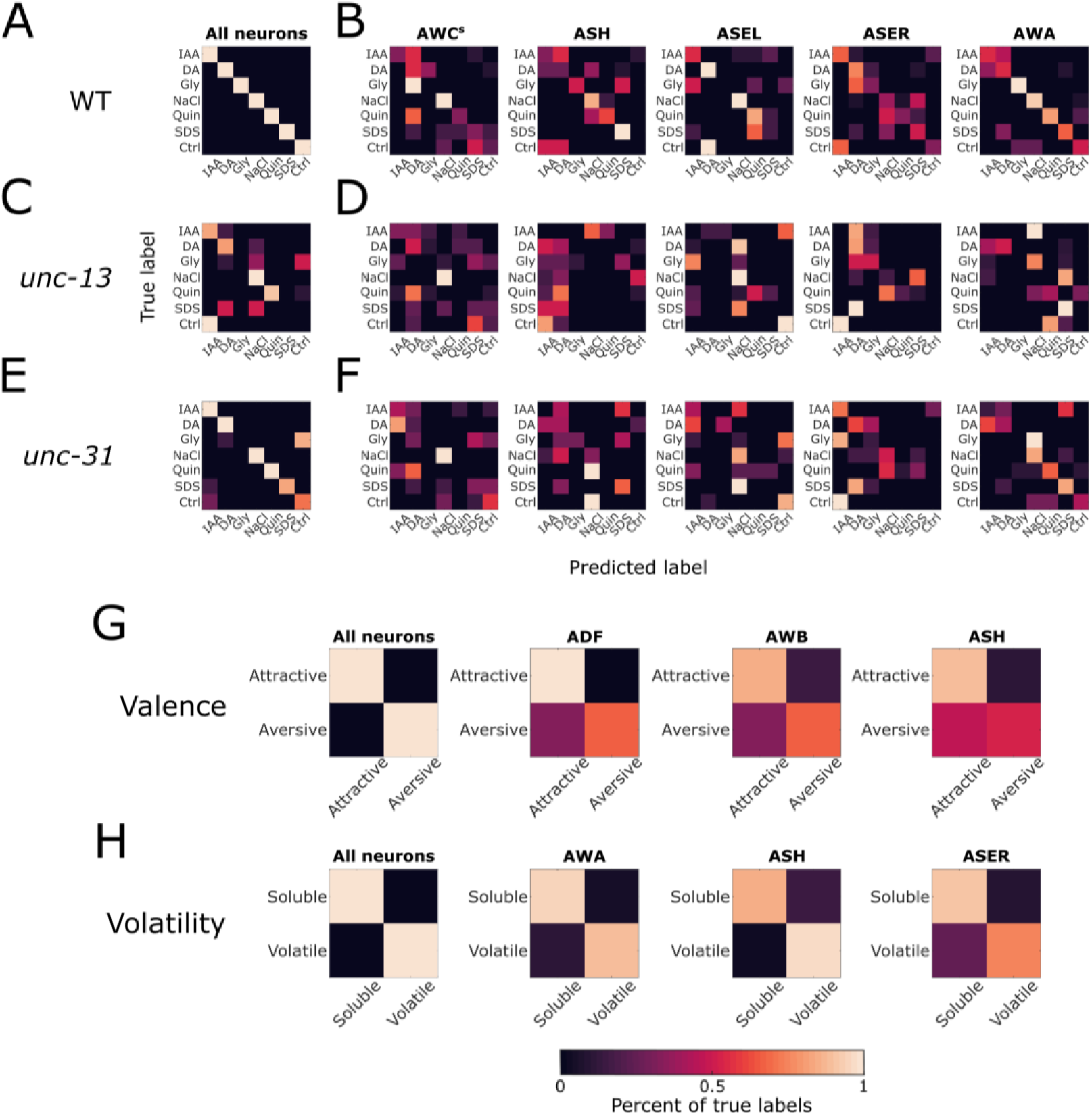
Neuronal activity predicts stimulus identity. Confusion matrices of a random forest classifier (100 trees and a depth of 4) trained on response dynamics of WT animals. The classifier was applied to test predictions on WT (**A, B**), *unc-13* (**C, D**), and *unc-31* (**E, F**) neuron activities. (**A, C, E**) Training data contained the responses of the entire network. (**B, D, F**) Training data contained single neuron responses to both ON and OFF steps of the stimuli. (**G-H**) Confusion matrices for classifying by volatility (**G**) and valence (**H**). The performance scale bar is the same for all panels.

Next, we analyzed the relative contribution of individual neurons to coding each of the stimuli by training the classifier on the ON and OFF responses of a single neuron. While the classification accuracy using individual neurons was generally low, neurons varied in the degree and type of classification they allowed for (**Figure 5B and supplementary figure 5**). For example, activity of the olfactory AWA neurons is sufficient to differentiate between the volatile IAA and DA and the other stimuli, however, as evidenced by the frequent mutual misidentification of the two, it is insufficient to distinguish between the two volatiles. Similarly, using the ASH neurons only, the classifier divided the stimuli into three distinct groups (IAA and DA, Gly and SDS, and NaCl and Quin), but had difficulties differentiating between the stimuli within each group. These results indicate that individual neurons contribute in varying degrees to the signaling of certain stimuli, but when combined, provide sufficient information to accurately identify all of the stimuli in our sample.

We next asked whether the internal communication (neurotransmission, neuropeptide release) in the network is crucial for stimulus identification. For this, we used the classifier trained on neuronal activity recorded from WT worms to predict stimuli based on the neuronal activity of the *unc-13* and *unc-31* mutants. Tested on *unc-31* data, the classifier performed nearly as well as on WT data, suggesting that neuro-peptidergic signaling plays a relatively minor role in stimulus identification (**Figure 5C-D**). In contrast, the classifier poorly predicted the stimuli in *unc-13* mutants, suggesting accurate coding is heavily reliant on neurotransmission (**Figure 5E-F**).

Prompted by the coarse separation of the classifier when relying on single neurons, we asked whether certain neurons are tuned towards specific properties of the stimulus. We therefore divided our stimulus sample by volatility (IAA and DA - volatile, NaCl, Gly, Quin, and SDS - nonvolatile) and valence (IAA, DA, and NaCl - attractive, Gly, Quin, and SDS - aversive), and trained the classifier again using the entire set of chemosensory neurons as well as with single neurons. The classifier performed well on both categories when trained on all chemosensory neurons (**Figure 5G-H**). Individual neurons varied widely in their ability to classify stimuli by category, where classification of volatility was best achieved using AWA, ASH, and ASE, whereas classification of valence was best when using the ADF, ASH, and AWB neurons (**Supplementary Figures 6-7**).

Together, our results suggest that individual neurons can encode specific features of a stimulus, and that precise stimulus identification is achieved when combining a small number of responding neurons. Moreover, neurotransmitter, rather than neuropeptide, signaling plays a pivotal role in modulating neural responses to allow stimulus discrimination.

### Temporal dynamics improves stimulus discrimination

This far, we showed that considering peak neural dynamics sufficed to accurately identify all stimuli in our data (**Figure 5A**). However, this could be due to the diverse nature and the small sample size of the stimuli used herein, as well as the large amount of neuronal data collected per trial. But could stimulus discrimination be improved by taking into account activity dynamics, in addition to peak activity?

To utilize time series in the classifier, we used CAnonical Time-series CHaracteristics to reduce the dimensionality of the response dynamics (Lubba et al. 2019). This reduced each neuron’s activity trace to 22 features that describe the time series (disregarding amplitude), including its linear and non-linear autocorrelation, scaling of fluctuations, distribution of values, etc. We then performed a principal component analysis on these features, and retrained the classifier by adding the first three components of each neuron (together explaining 55% of the variance, **Supplementary figure s8**) as variables. Thus, each dataset consisted of a single neuron’s response represented by eight variables – ON and OFF step response magnitudes, and a total of six trace features. We next compared the performance (F1-score from cross-validation) of the classifiers when trained on the response amplitude only, on the three principal components of the trace dynamics only, and the combined amplitudes and trace dynamics for each individual neuron (**Figure 6A-D**).

**Figure 6.**
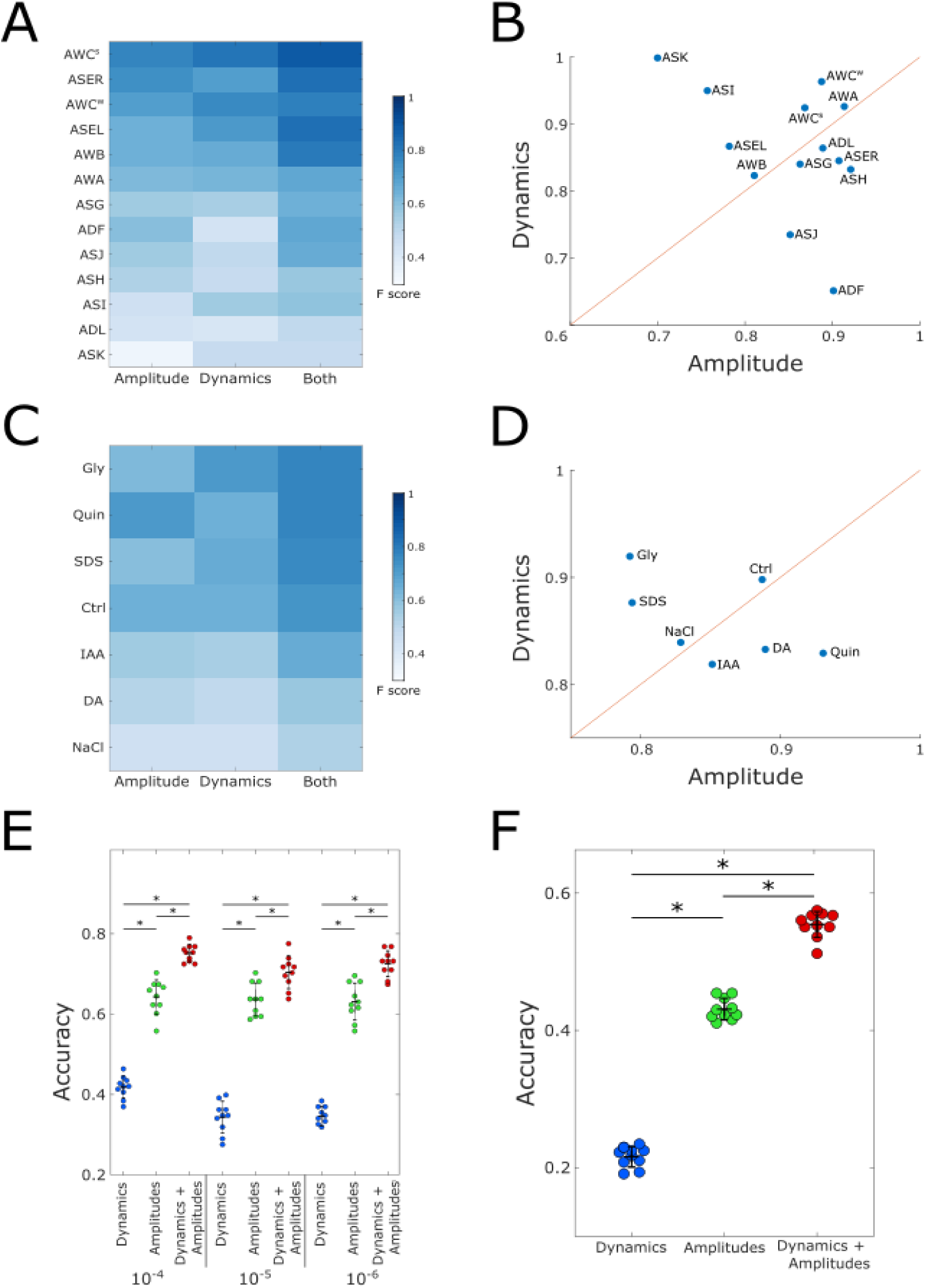
Temporal dynamics provides additional information for stimulus identification. **(A)** Global F1-scores (across all conditions) for each neuron when considering only the trace dynamics, the amplitudes, or both. **(B)** Scatter plot depicting the contribution of the amplitudes and the response dynamics to the overall performance of classification by each neuron as shown in **(A)**. Amplitudes and dynamics are expressed as a fraction (relative contribution) of their combined accuracy as shown in the third column of panel **(A)**. **(C)** Averaged F1 scores across all neurons for each condition when considering only the trace dynamics, the amplitudes, or both. **(D)** Scatter plot depicting the contribution of the amplitudes and the dynamics to the overall Performance of classification of each stimulus as shown in **(C)**. Amplitudes and dynamics are expressed as a fraction (relative contribution) out of their combined accuracy as shown in the third column of panel (C). **(E)** Classifier accuracy scores predicting stimulus identity based on dynamics, amplitudes, and both. The data used herein was obtained from (Lin et al. 2023) consisting of 11 sensory neurons across 23 different stimuli at 10^-4^, 10^-5^ and 10^-6^ concentrations. *p< 10^-3^ (one sided t-test, FDR corrected). **(F)** Classifier accuracy scores predicting stimulus identity based on dynamics, amplitudes, and both when combining all the data from Lin et al (irrespective of the specific concentration). *p< 10^-4^ (one sided t-test, FDR corrected).

For most neurons, considering both aspects of the response had an additive effect, where performance of the classifier trained on both trace dynamics and amplitudes was better than each by itself (**Figure 6A**). However, a classifier that was trained on only the dynamics features of either AWCW or ASI, performed better than when trained on amplitudes alone, and combining traces and amplitudes of these neurons resulted in minimal additional improvement (**Figure 6A-B**). In contrast, for ASJ and ADF, the amplitude provided the most information, and taking trace features into account did not improve performance further.

We also estimated the contribution of amplitudes and trace dynamics to the representation of each stimulus by taking the mean F1-score of each stimulus over all neurons (**Figure 6C-D**). For most stimuli, the effect of combining amplitudes and trace features was additive. Interestingly, overall performance of the classifier tended to be better for the aversive stimuli (Quin, SDS, and Gly) than for the attractive ones (IAA, DA, NaCl), suggesting that the sensory system is more finely tuned to precisely identify noxious stimuli (**Figure 6C)**.

These results suggest that given a sufficiently diverse stimulus space, the identities of the stimuli can be efficiently encoded using response amplitudes alone (**Figure 3A and figure 5A**). However, response dynamics of individual neurons carry considerable additional information that could be used to help distinguish between more closely related stimuli, particularly aversive ones.

To better understand the relative contribution of response dynamics to stimulus coding, we analyzed available data that measured the activity of all amphid neurons in response to a panel of 23 different odorants, spanning six chemical classes, each in several concentrations (Lin et al. 2023). We first extracted peak activities and repeated the analysis with our classifier.

Overall, our classifier performed similarly to the one described in the paper, reaching comparable prediction accuracy of ∼70% when trained on response amplitudes alone (**Figure 6E**). Training the data using the dynamics yielded a lower accuracy of ∼40% for each individual concentration. However, combining amplitudes with response dynamics significantly improved the performance of the classifier (for each of the dilutions), thus mirroring the single-neuron classification results obtained in our data (**Figure 6A-D**).

We then applied the classifier to the entire dataset (23 odorants at three concentrations, for a total of 69 individual stimuli). Due to the similarity of the population coding to different concentrations of the same stimulus (Lin et al. 2023), it should be particularly challenging to tell them apart. However, even with such a large number of stimuli, the dynamics significantly added to the overall accuracy compared to the performance when considering amplitudes alone (**Figure 6F**).

Taken together, these findings indicate that combining response amplitudes with response dynamics significantly improves stimulus identification by generating unique codes to each stimulus (and its concentration), effectively enhancing the coding capacity of the chemical space across a wide range of concentrations.

## Discussion

We studied how a compact chemosensory system, consisting of limited neural resources, encodes a variety of chemical cues. Using *C. elegans* as a model system, we found that animals use a small set of neurons to encode each of the stimuli, where ∼2-4 sensory neurons are the primary sensors of the stimulus. A few additional sensory neurons are recruited via inter-neuronal signaling. Interestingly, while most neurons show a stereotypic response dynamics of sharp increase in calcium levels followed by a slow decay, some neurons show variable dynamics that depend on either stimulus identity or inter-neuronal communication. These fine response dynamics significantly improve stimulus identification, effectively enhancing the coding capacity of the compact sensory system.

To analyze a wide space of possible sensory responses, we employed a variety of stimuli that represent both olfactory and gustatory cues, some of which are attractive while others are repulsive (**Figure 1**). Overall, our results agreed with previous works in regards to response profiles of the sensory system to various chemical stimuli (Suzuki et al. 2008; Leinwand and Chalasani 2013; Leinwand et al. 2015; Chalasani et al. 2007; Zaslaver et al. 2015; Yemini et al. 2021; Lin et al. 2023). In addition, most symmetric neuron pairs showed highly correlated activity, with the only exceptions being the AWC and the ASE neuron pairs (**Figure 2**). However, it is plausible that testing additional stimuli will reveal more neurons with differential bilateral functionality.

Our results recapitulate a previous report showing that the neurons respond in a hierarchical manner, where some are broadly tuned (*e.g.,* AWC) to respond to all stimuli, whereas others are more finely tuned and respond to specific stimuli only (Zaslaver et al. 2015). A similar principle was also observed when analyzing responses to a wide array of olfactory stimuli (Lin et al. 2023), though some of the broadly and narrowly tuned neurons differed between this and our study. This difference could be due to the nature of the stimuli used in each study. For example, the ASI neurons were broadly activated in our study mostly in response to gustatory stimuli. This may explain why this neuron was not detected as a broad responder when assaying responses to volatile cues (Lin et al. 2023). Thus, the segregation to broadly- and narrowly-tuned neurons may heavily depend on the stimuli used. Nevertheless, in both studies, AWC is consistently identified as a broadly responding neuron.

While a distinct set of neurons encodes each of the stimuli, the response to each stimulus was sparse, in line with previous findings (Zaslaver et al. 2015). Analyses of mutant strains revealed that neurotransmission is the main route in which primary neural responders recruit and activate secondary neurons (**Figure 4 and supplementary figure 3**). In contrast, neuropeptides primarily modulate the activity of the responding ensemble. Additional intern-neural signaling route involves electrical gap junctions. For example, ASH responses are regulated non-cell-autonomously by gap junctions (Krzyzanowski et al. 2013). *C. elegans* contains 25 genes coding for innexins (Altun et al. 2009; Simonsen, Moerman, and Naus 2014; Jin et al. 2020), the invertebrate analogs of the vertebrate connexins that make up electrical gap junctions. This substantially complicates analysis of their involvement in recruitment of secondary responders. Nevertheless, if gap junctions also play a role in activating secondary neural responders, then the actual number of primary neurons might be lower than the 2-4 neurons based on our findings (**figure 4B**). This possibility further lowers the already limited coding capacity of the sensory system and underscores the importance of inter-neural communication to fine tune response dynamics that improve stimulus identification.

While we find a small fraction of the chemosensory neurons to respond to each of the stimuli, Lin et al found that a higher fraction of the sensory neurons is activated when presented with volatiles only (Lin et al. 2023). The lower number of responding neurons in our analyses may be because we separated the responses to ON and OFF steps since some of the neurons are classically on-step responders (e.g., AWA) while others are consistently off-step responders (e.g., AWC). Furthermore, we considered the two bilateral AWC and ASE neurons as distinct entities, analysis that further contributed to the sparser coding conclusion.

The sparse representation in the chemosensory system presents a combinatorial problem for the animal, where potentially thousands of different chemical cues are encoded by a small subset of the sensory neurons. This problem is exacerbated by the further division of responding neurons into primary (directly sensing) and secondary (network recruited) neurons, as revealed by the inter-neuronal signaling mutants. This observation bears some resemblance to the “primacy coding” model proposed for mammalian olfactory systems, whereby the identity of a stimulus is encoded by a small set of early-responding glomeruli in a concentration-independent manner (Wilson et al. 2017; Chong et al. 2020). According to this model, coarse stimulus identification occurs first, with fine-tuning of odor identity and concentration being mediated by higher-latency glomeruli. While the relatively slow Calcium dynamics does not allow us to resolve response latency, it is an intriguing possibility that a similar “primacy code” occurs in *C. elegans*, with the broad categorization of the stimulus being determined by primary responding neurons, followed by secondary-responder mediated fine-tuning.

Furthermore, our analyses showed that the relative contribution of the response dynamics to stimulus identification differs between neurons (**Figure 6**). It is appealing to consider this feature as another variation of a “primacy code”, where the initial response amplitude provides the coarse stimulus identification, and the longer-scale temporal dynamics of the same neuron assists the secondary responders in its fine-tuning. Notably, these temporal features may be particularly important in *C. elegans* as they can propagate from the sensory neurons across the network to direct matching behavioral outputs (Kato et al. 2014; Itskovits et al. 2018).

Activity of secondary responding sensory neurons relies on lateral signaling within the sensory layer or through feedback from downstream interneurons. The importance of such signaling was demonstrated, for example, in the improvement of the signal-to-noise ratio to support a more robust chemotaxis (Chalasani et al. 2007; 2010). Recruitment of unique subsets of secondary neurons could also increase coding capacity, potentially alleviating the combinatorial limitations of a small sensory system and the limited number of primary responding neurons. For example, if two different stimuli activate the same primary responding neurons, the secondary recruited neurons may differ. This could be due to the fact that each sensory neuron expresses several chemical receptors (Hammarlund et al. 2018; Taylor et al. 2021). Thus, even if two different stimuli elicit a response in the same sensory neurons, distinct intracellular signaling paths may lead to two different synaptic outputs that will eventually recruit different secondary responders. This in turn generates stimulus-specific codes that allow the animal to discriminate between the two stimuli, despite the fact that they are sensed by the same sensory neurons.

We propose an additional strategy by which the coding capacity could be increased. Our results show that the sensory neurons exhibit a variety of possible calcium dynamics determined by both extrinsic (stimulus) and intrinsic (inter-neuronal signaling) factors. Attempting to predict stimulus identity based on single-neuron responses revealed that these dynamics carry additional information that may help differentiate between stimuli. This may be particularly useful when confronted with a large stimulus space composed of many closely-related compounds, or different concentrations of the same stimulus.

*C. elegans* neurons have long been thought to only employ graded potentials. However, calcium-mediated action potentials have been described in several neurons, including the AWA sensory neurons (Lin et al. 2023). In AWA, calcium transients correlate with spike trains suggesting that different activation frequencies may code for different stimuli, thus possibly further increasing the coding capacity.

Together, this study reveals the principles by which a chemosensory system that is limited in its sensory resources may uniquely encode a large repertoire of chemical stimuli. Similar strategies may have evolved in higher brain systems where a more elaborate sensory system is required to uniquely code and discriminate between a greater space of different stimuli.

## Materials and Methods

### Strains

ZAS280 *In*[*osm-6*::GCaMP3, *osm-6*::ceNLS-mCherry-2xSV40NLS] (Iwanir et al. 2019).

ZAS325 is a cross between ZAS280 and *unc-31*(e928) (Pritz et al. 2023).

ZAS371 is a cross between ZAS280 and *unc-13*(s69).

### Worm cultivation

All worms were grown on NGM plates seeded with OP 50 and kept at 20 °C. Age synchronization was performed by bleaching. All experiments were done 3 days post the bleach, using young adult worms.

### Imaging the chemosensory system

Worms were starved for 20 minutes on un-seeded NGM plates, and then inserted into a microfluidic chip (Chronis, Zimmer, and Bargmann 2007) where they were partially anesthetized with 10mM levamisole and left to habituate for 10 more minutes. Recordings lasted for 2.5 minutes starting with 30 seconds of light habituation. ON/OFF steps lasted for 1 minute each, with stimulus presentations occurring at 30 and stimulus removal at 90 seconds. Since ASH and ASK showed strong responses to light itself, for the ON step responses of these neurons, we used recordings collected from a second round of ON steps.

Stimuli used were diacetyl 10^-4^ (DA), isoamyl alcohol 10^-4^ (IAA), NaCl 50 mM (NaCl), glycerol 1 M (Gly), SDS 0.1% (SDS), and quinine 5mM (Quin).

Imaging was performed on a Nikon A1R+ confocal laser scanning microscope with a water immersion x40 (1.15NA) objective at ∼1.5 volumes/second and with Z-axis intervals of 0.5-0.8 µm. To account for slightly varying acquisition rates, all traces were interpolated to 2Hz. The system was controlled by the Nikon NIS-elements software.

### Neuron identification and signal extraction

Neurons were detected and tracked based on the nuclear mCherry signal using an algorithm developed by (Toyoshima et al. 2016). Neuronal identities were determined visually based on their anatomical positions. Neurons that could not be unambiguously identified were removed from analysis. The left-right assignment of the AWC^ON^/AWC^OFF^ neurons is random in each worm (Troemel, Sagasti, and Bargmann 1999), and our reporter strain does not allow to differentiate between them. To maintain the functional distinction, the AWC pair in each worm was sorted by activation strength. Where relevant, the neurons are marked AWC^s^ (strong) and AWC^w^ (weak).

Calcium traces were extracted from within 0.9 of the radius of the originally segmented neural sphere, in an effort to reduce cross-reads from nearby neurons. In cases where cross-reads persisted, the signal was read from the half of the sphere facing away from the signal donor. Signal intensity was normalized by baseline activity which was defined as the lowest value of a 10-frame running average. All data analyses were performed using in-house Matlab, Python, and R scripts.

### Time-trace correlations

To calculate the correlations in neuron dynamics, pairwise correlations were calculated for all neurons in each worm, and then averaged across all worms in a given condition. Overall correlations were determined by averaging all of the worms in all conditions. Due to the strong correlations between bi-laterally symmetrical neurons, in the statistical analysis of all neurons (excluding the AWC and ASE pairs), the left and right sides were pooled together, unless specified otherwise. The AWC neurons within each worm were sorted by overall activity levels and designated AWC^s^ (strong) and AWC^w^ (weak).

### Calculating response magnitudes

To determine response magnitudes, average baseline activity of 10 seconds prior to the step was subtracted from the maximum activity in the 7 seconds following the step. To account for activity caused by changes in the flow direction and the fluorescent dye, the responses of the control group were compared to zero using one-sample t-test, and all subsequent WT conditions were compared to the control group using two-sample t-test. Responses were deemed significant if they were significantly different from the control, regardless of the response of the control. Mutant responses were compared to the WT response to the same stimulus. Multiple comparisons correction was performed using FDR.

For classification, missing data was imputed as described in (Lin et al. 2023), using a matrix completion algorithm based on minimization of the nuclear norm provided by (J. Fan, L. Ding, Y. Chen, M. Udell 2019). (https://github.com/udellgroup/Codes-of-FGSR-for-effecient-low-rank-matrix-recovery).

### Neuron activity dynamics

For trace dynamics analysis, only the neurons that were determined to respond (and therefore displayed response dynamics to analyze) were used. The response traces of each step were aligned to the maximum first derivative value around the stimulus presentation, and z-normalized. Each trace lasted from 20 seconds before the stimulus presentation to 50 seconds after. PCA was performed on the individual traces, and the mean trace was calculated per neuron, per stimulus, and projected onto the PC space. Clustering was performed using K-means.

### Stimulus identity classification

A random forest classifier (with 100 trees and a depth of 4) was used to predict the identity of a stimulus from neuronal responses. ON and OFF step responses of all neurons from each worm were concatenated into 26-dimensional vectors, which were then randomly divided into training (80%) and test (20%) sets. For single-neuron classification, each point consisted of the ON and OFF step responses of one neuron. For cross-validation, the model was trained 10 times on different train-test sets, and the average F-score across all 10 trials was calculated. The contribution of response amplitudes and trace features to classification by neuron or stimulus was calculated by taking the mean F-score across stimuli/neurons respectively. For example, the success of the classification of diacetyl was calculated as the mean F-score of diacetyl across all individual neurons. Similarly, success of classification by AWA was calculated as the mean F-score of AWA across all stimuli.

### Classification by dynamics

For the purpose of classification, the dimensionality of neuronal activity dynamics was reduced to 22 trace features using the Catch22 algorithm (Lubba et al. 2019). These features were then z-normalized and PCA was performed. The first 3 principal components were used as additional features for the random forest classifier. (https://github.com/DynamicsAndNeuralSystems/catch22).

### Resampling data

The data provided in (Lin et al. 2023) contain neuron activity traces grouped by the stimulus presented. Our classifier, however, requires the data to be arranged as “complete” worms, where each observation (worm) consists of the activity features of all of the amphid neurons. These “synthetic” worms were constructed by independently and randomly sampling each neuron from all of the responses of that specific neuron to a specific stimulus. E.g. a synthetic diacetyl-sensing worm was composed of a random AWA response to diacetyl, an independently randomly chosen AWB response to diacetyl, etc. Where trace features were also used, all features of a neuron were sampled from the same response trace (i.e. the amplitude and trace feature PCs were taken together, rather than each being independently sampled). The model was cross-validated 10 times with different train-test (80%/20%) splits and the mean F-score was calculated.

## Data availability

All the data and the scripts that analyzed the data to produce the results and the figures herein are available through Github: https://github.com/zaslab/Sensory-encoding

---

**Supplementary figure 1.**
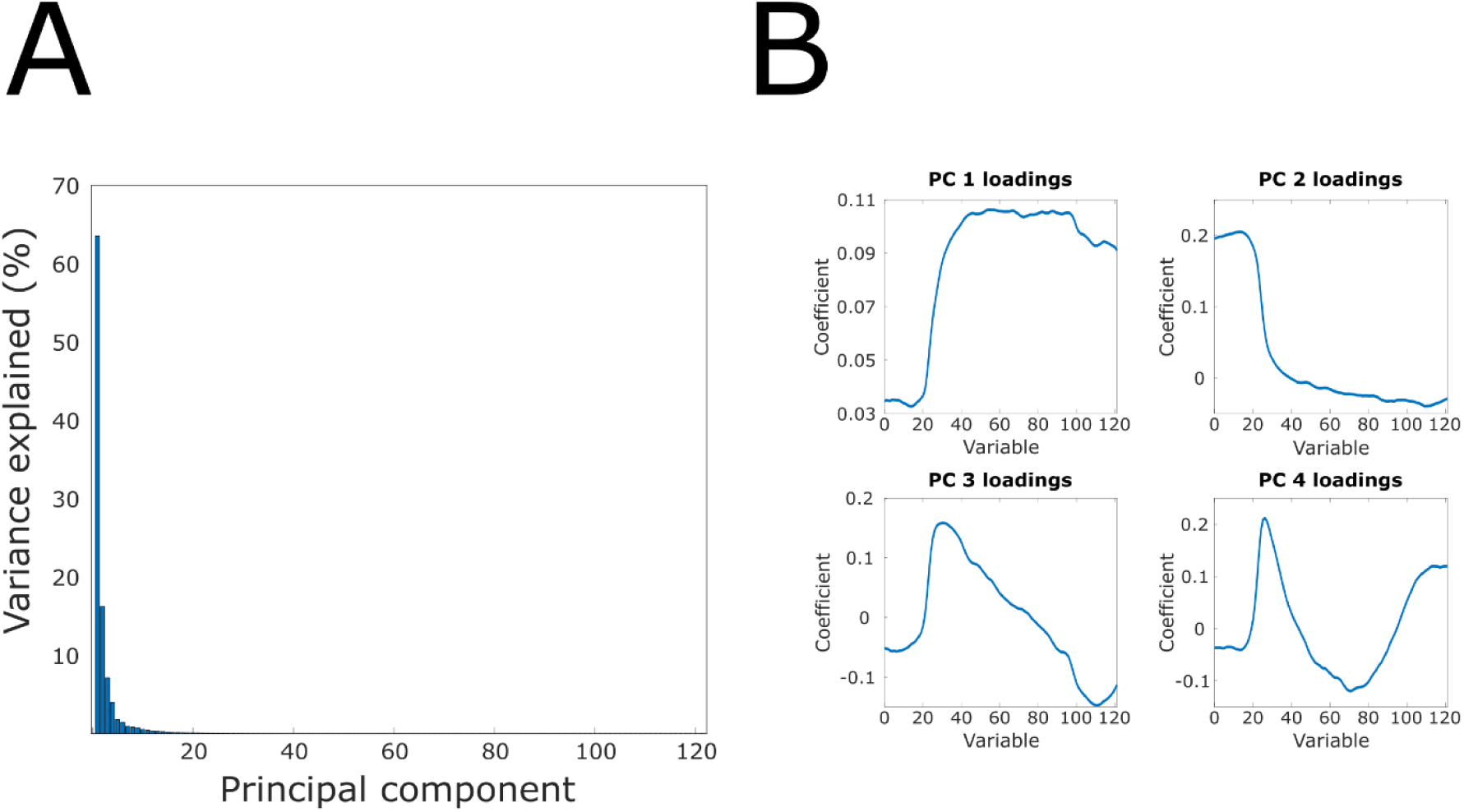
Principal component analysis of the chemosensory response dynamics. **(A)** Percent of variance explained by each of the principal components. **(B)** Time-point PC loadings of the first four principal components.

**Supplementary figure 2.**
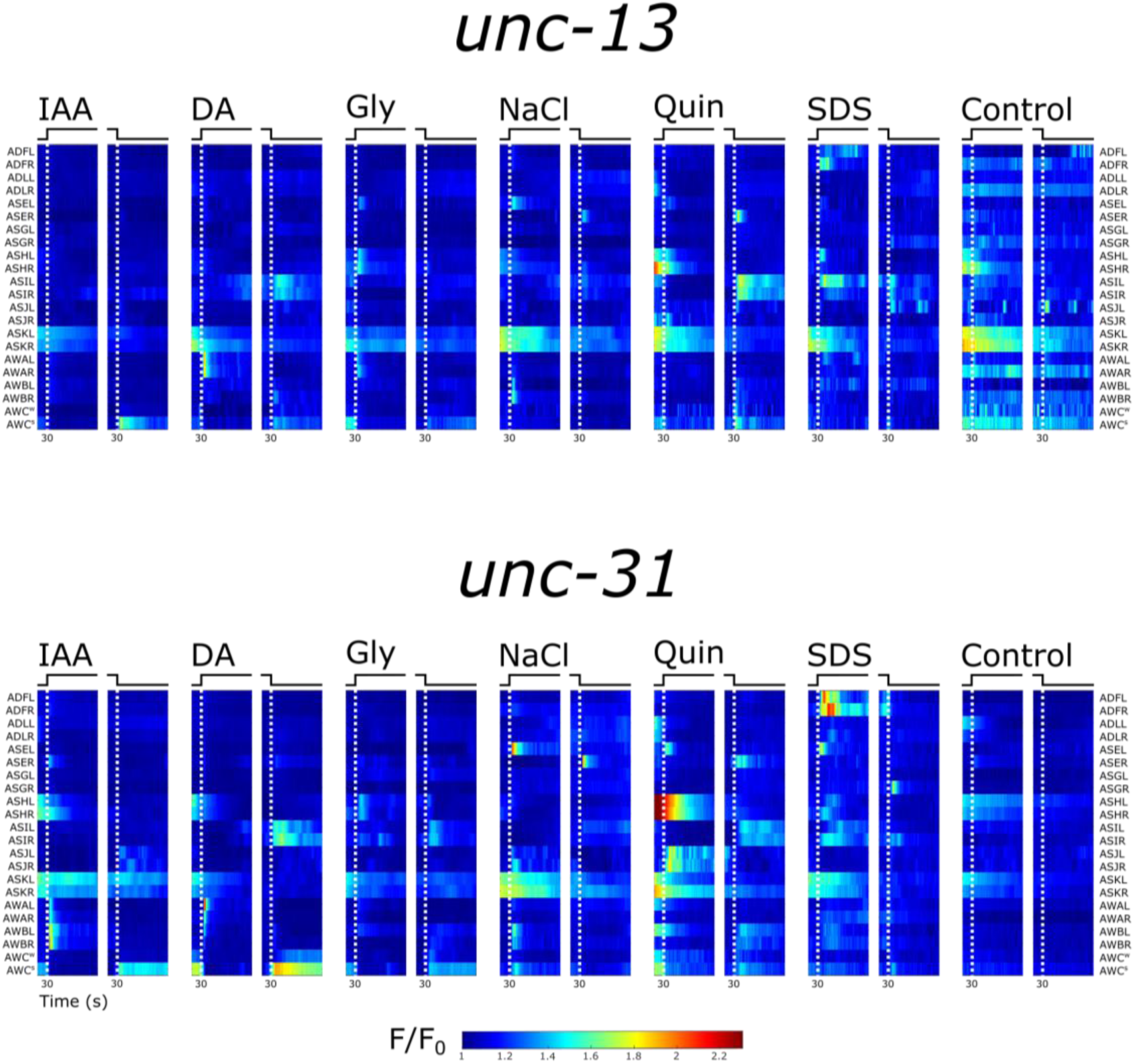
Functional dynamics of the chemosensory neurons in the *unc-13* and *unc-31* mutant strains. Mean neural dynamics of individual neurons following stimulus presentation and removal. White dashed lines indicate ON/OFF steps. Conditions tested: Control; DA, diacetyl 10^-4^; IAA, isoamyl alcohol 10^-4^; NaCl, sodium chloride 50 mM; Gly, Glycerol 1M; Quin, Quinine 5 mM; SDS 0.1%. A fluorescent red dye (500 nM rhodamine) was added to the stimuli to verify accurate stimulus switch. The control condition consisted of switching between buffer and buffer+dye. Responses observed in the control condition served as the baseline responses for neurons that may have responded to the dye only. The AWC pair is sorted by activation strength in each worm and is marked AWC^s^ (strong) and AWC^w^ (weak). Note that the ASK, ASH and ADL neurons respond to blue light, hence the activity at the start of the imaging period.

**Supplementary figure 3.**
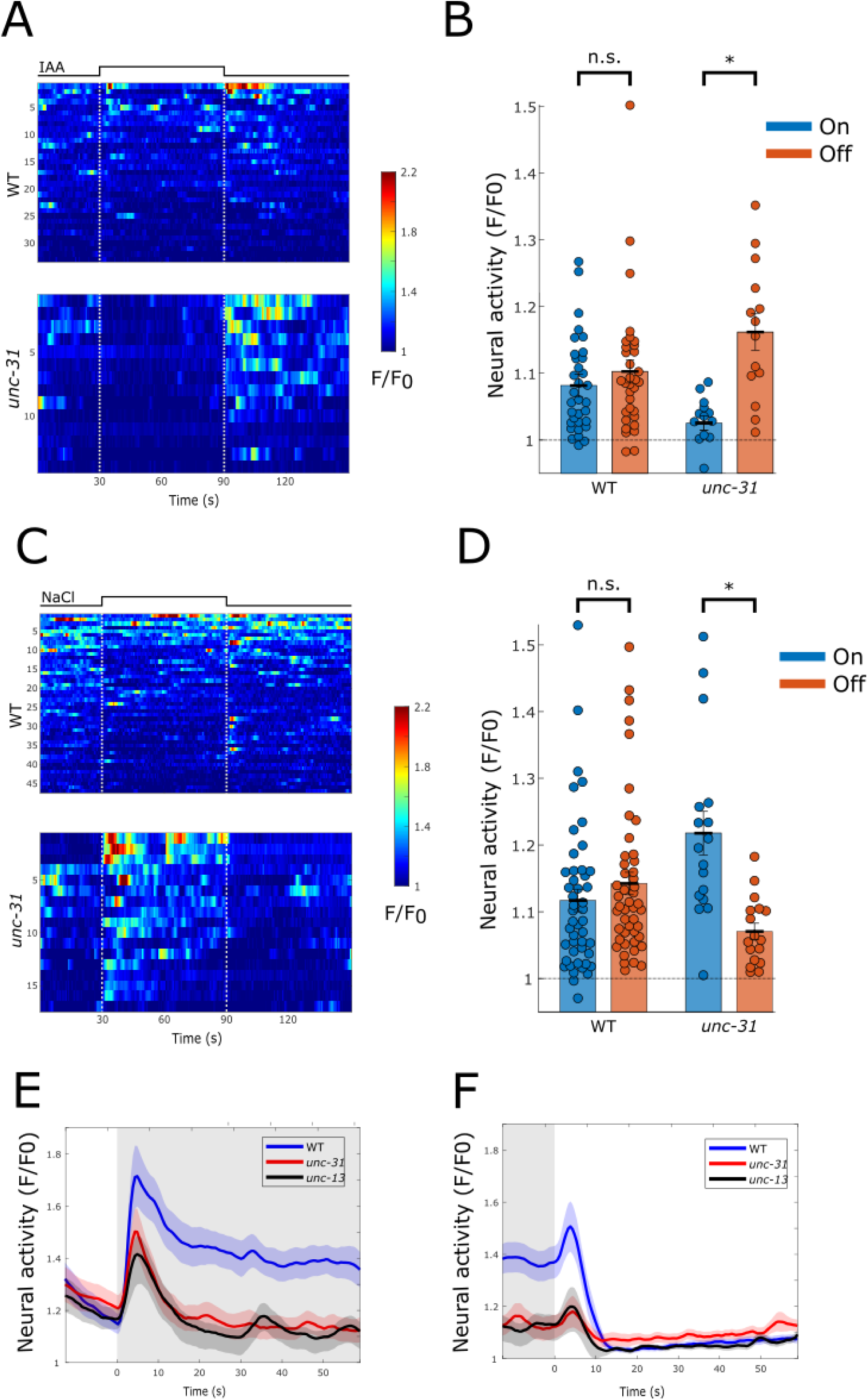
Modulated activity is stimulus specific and also depends on inter-neural signaling communication. **(A-D)** ASJ activity is modulated by neuropeptides and is also stimulus specific. **(A,B)** ASJ responses to the stimulus IAA. **(C,D)** ASJ response to the stimulus NaCl. **(A,C)** Each line represents an individual ASJ neuron throughout the entire imaging period. Top, responses of WT worms. Bottom, responses of *unc-31* mutants. Each row represents an individual ASJ neuron throughout the entire imaging period. Dotted white lines indicate the ON (t=30 s) and the OFF (t=90 s) steps. **(B,D)** Bar graphs showing the mean activity level over the entire length of the ON/OFF steps. Two-sided t-test, *\*p* < 0.01. The differential responses between WT and the *unc-31* mutant animals suggest that neuropeptide signaling affects ASJ activity. The differential activity of WT animals in response to IAA and NaCl suggests that ASJ activity is also stimulus specific. **(E, F)** Activity dynamics of the ASH neurons in response to **(E)** ON and **(F)** OFF steps of 1M glycerol in WT and mutant worms (*unc-31* and *unc-13*). In both cases, it is evident that ASH responses are modulated by both neuropeptide and neurotransmitter release.

**Supplementary figure 4.**
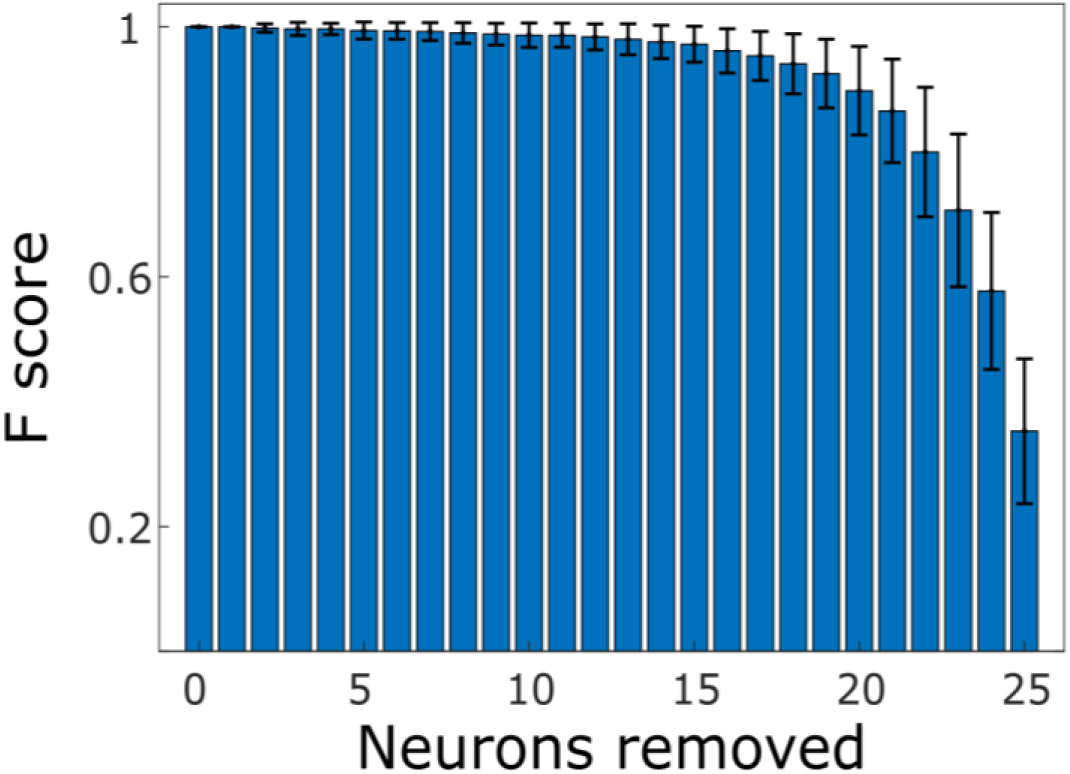
Classifier performance decreases the more neurons are removed from the training set. Mean F score of the classifier trained with a sample of randomly removed sets of neurons. Error bars represent std.

**Supplementary figure 5.**
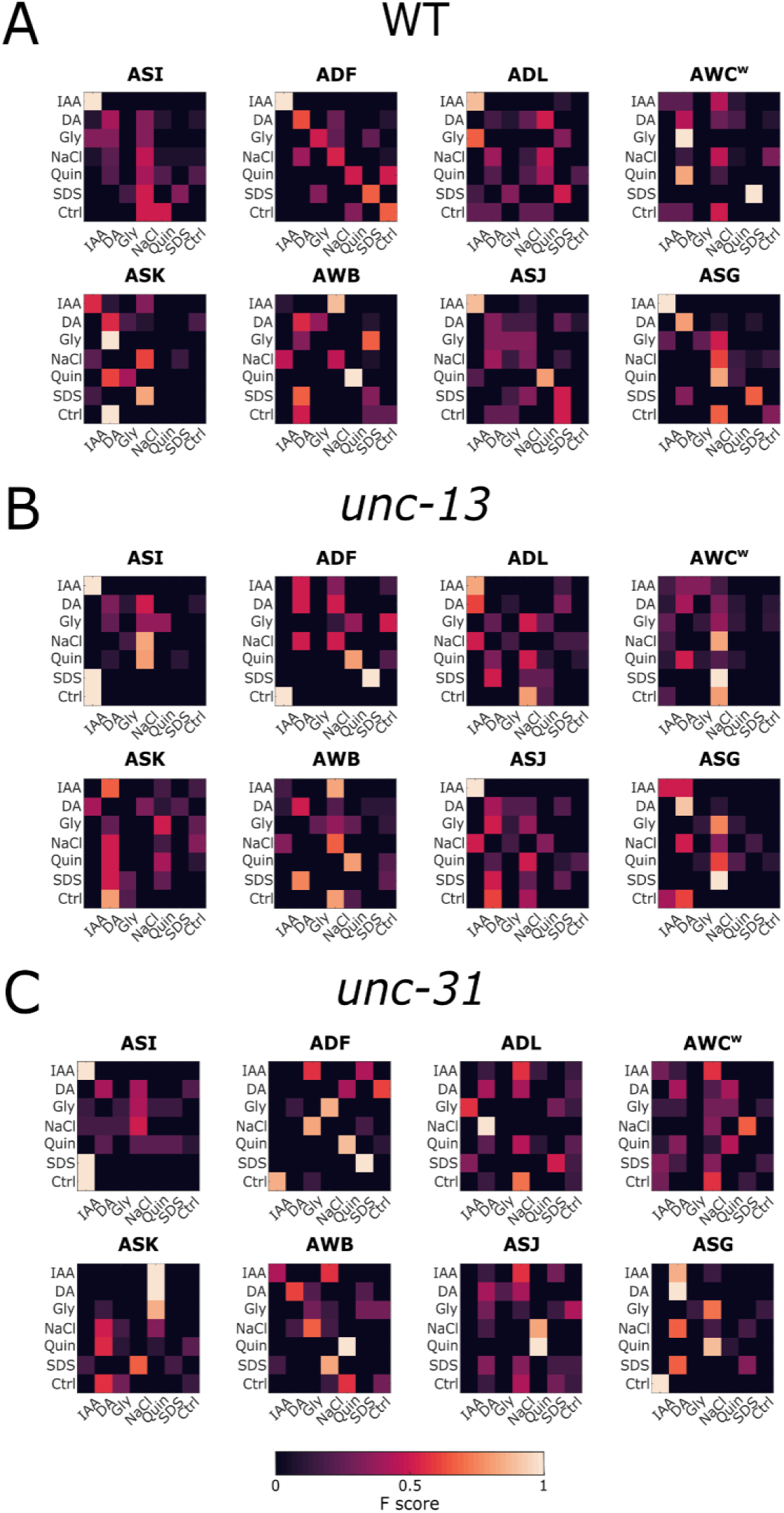
Neuronal activity predicts stimulus identity. Confusion matrices of a random forest classifier trained on single neuron responses (ON and OFF steps) of WT animals. The classifier was applied to test predictions on WT (**A**), *unc-13* (**B**), and *unc-31* (**C**) neuron activities.

**Supplementary figure 6.**
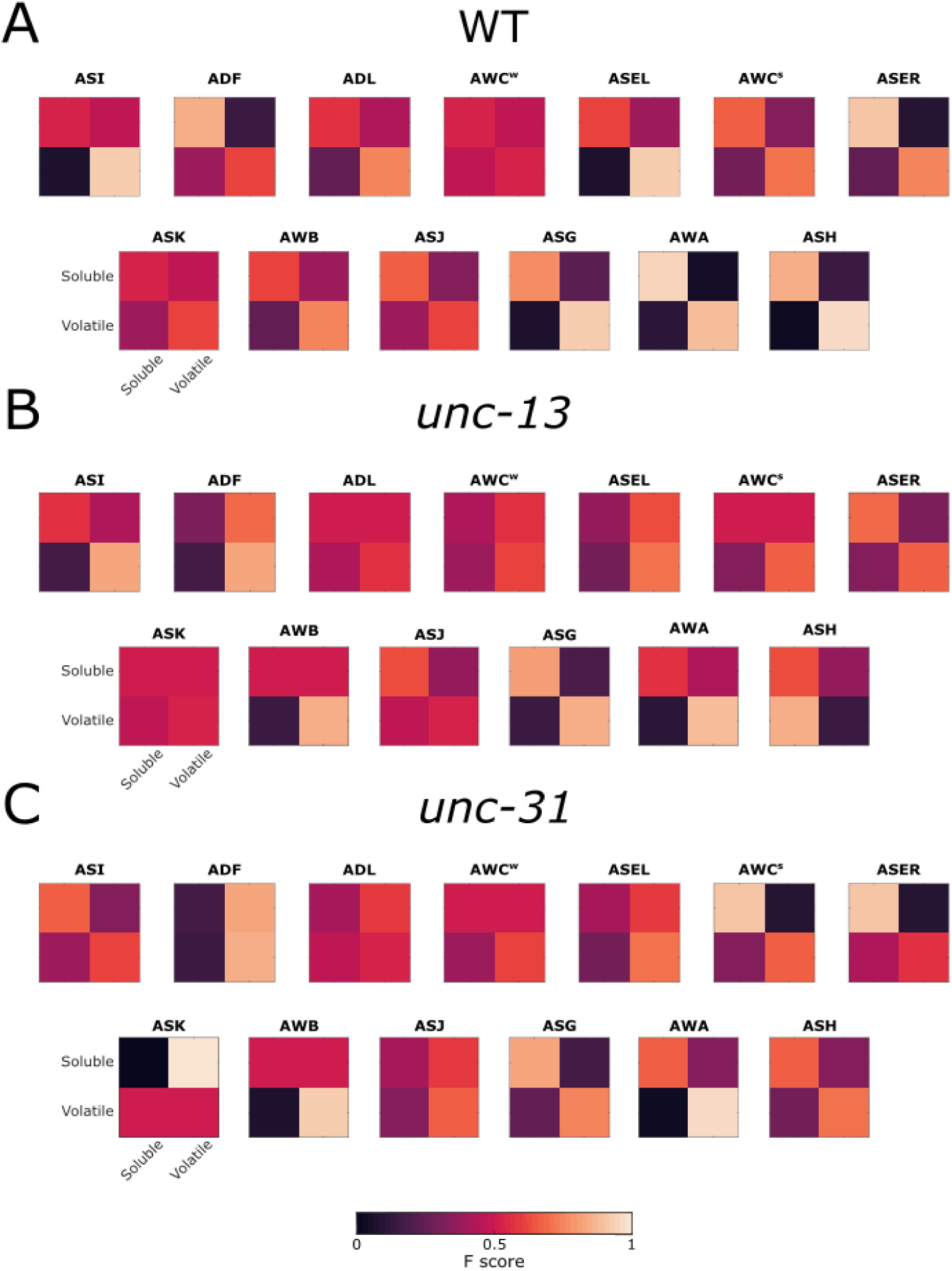
Neurons vary in their contribution to the prediction of the solubility of a stimulus. Confusion matrices of a random forest classifier trained to predict the solubility of the stimulus based on single neuron responses (ON and OFF steps) of WT animals. The classifier was applied to test predictions on WT (**A**), *unc-13* (**B**), and *unc-31* (**C**) neuron activities.

**Supplementary figure 7.**
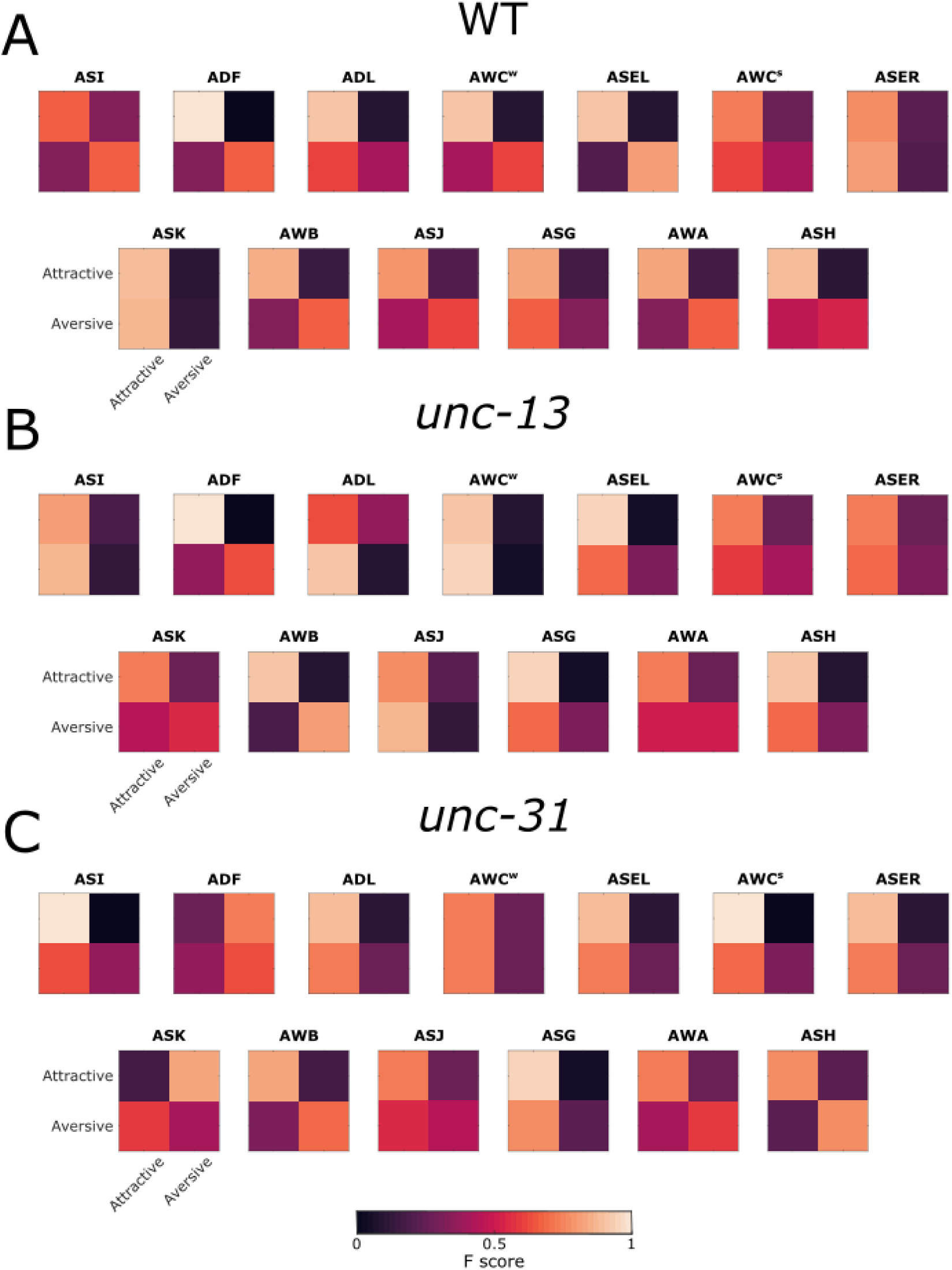
Neurons vary in their contribution to the prediction of the stimulus valence. Confusion matrices of a random forest classifier trained to predict the valence of the stimulus based on single neuron responses (ON and OFF steps) of WT animals. The classifier was applied to test predictions on WT (**A**), *unc-13* (**B**), and *unc-31* (**C**) neuron activities.

**Supplementary figure 8.**
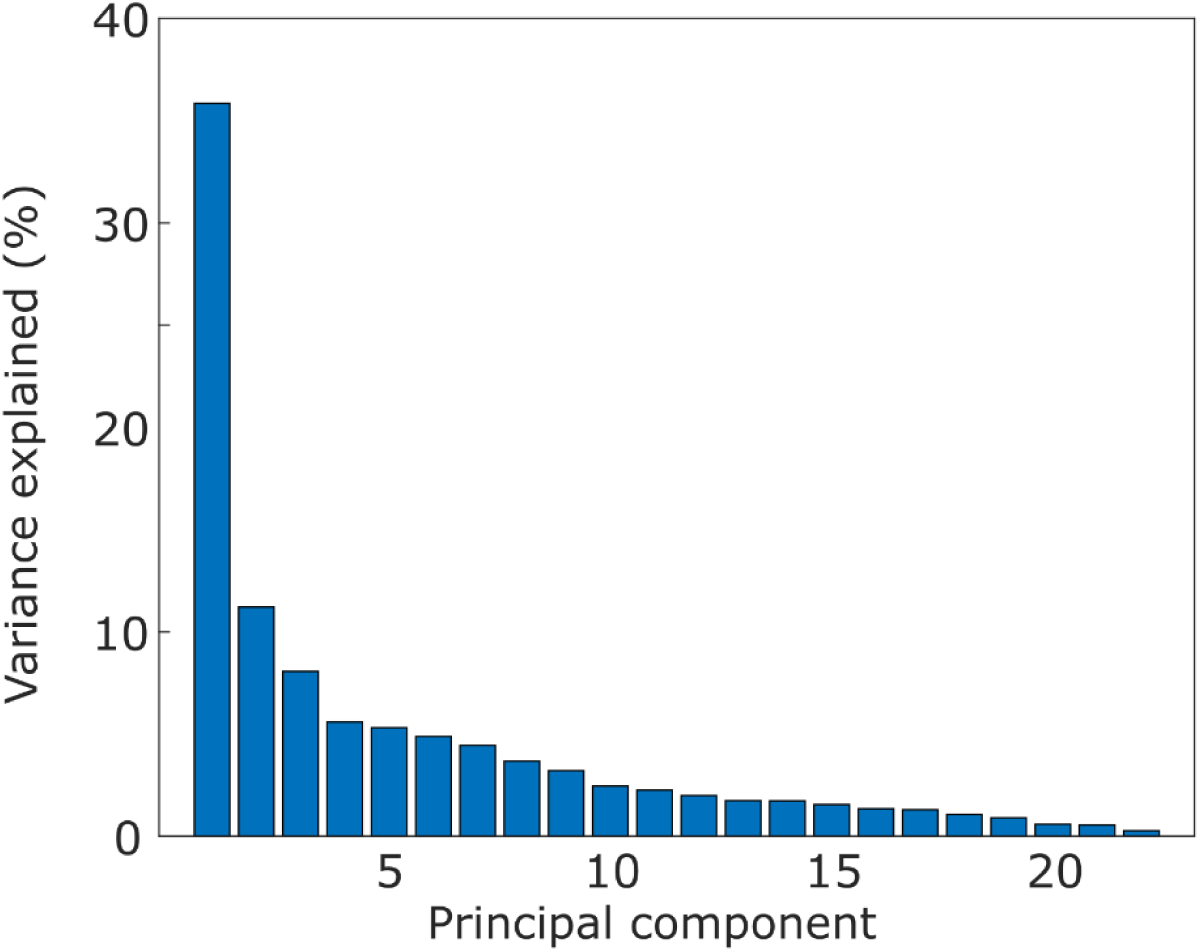
Variance explained by each principal component of the PC analysis performed on the trace features of neuronal responses extracted using the Catch22 algorithm (see Methods). The first three principal components explain ∼55% of the variance.

## References

Altun, Zeynep F., Bojun Chen, Zhao-Weng Wang, and David H. Hall. 2009. “High Resolution Map of Caenorhabditis Elegans Gap Junction Proteins.” Developmental Dynamics: An Official Publication of the American Association of Anatomists 238 (8): 1936–50. 10.1002/dvdy.22025.

Bargmann, Cornelia. 2006. “Chemosensation in C. Elegans.” WormBook. 10.1895/wormbook.1.123.1.

Chalasani, Sreekanth H., Nikos Chronis, Makoto Tsunozaki, Jesse M. Gray, Daniel Ramot, Miriam B. Goodman, and Cornelia I. Bargmann. 2007. “Dissecting a Circuit for Olfactory Behaviour in Caenorhabditis Elegans.” Nature 450 (7166): 63–70. 10.1038/nature06292.

Chalasani, Sreekanth H., Saul Kato, Dirk R. Albrecht, Takao Nakagawa, L. F. Abbott, and Cornelia I. Bargmann. 2010. “Neuropeptide Feedback Modifies Odor-Evoked Dynamics in C. Elegans Olfactory Neurons.” Nature Neuroscience 13 (5): 615–21. 10.1038/nn.2526.

Chong, Edmund, Monica Moroni, Christopher Wilson, Shy Shoham, Stefano Panzeri, and Dmitry Rinberg. 2020. “Manipulating Synthetic Optogenetic Odors Reveals the Coding Logic of Olfactory Perception.” Science 368 (6497): eaba2357. 10.1126/science.aba2357.

Chronis, Nikos, Manuel Zimmer, and Cornelia I. Bargmann. 2007. “Microfluidics for in Vivo Imaging of Neuronal and Behavioral Activity in Caenorhabditis Elegans.” Nature Methods 4 (9): 727–31. 10.1038/nmeth1075.

Cook, Steven J., Travis A. Jarrell, Christopher A. Brittin, Yi Wang, Adam E. Bloniarz, Maksim A. Yakovlev, Ken C. Q. Nguyen, et al. 2019. “Whole-Animal Connectomes of Both Caenorhabditis Elegans Sexes.” Nature 571 (7763): 63–71. 10.1038/s41586-019-1352-7.

Ferkey, Denise M, Piali Sengupta, and Noelle D L’Etoile. 2021. “Chemosensory Signal Transduction in Caenorhabditis Elegans.” Genetics 217 (3). 10.1093/genetics/iyab004.

Ghosh, D. Dipon, Michael N. Nitabach, Yun Zhang, and Gareth Harris. 2017. “Multisensory Integration in C. Elegans.” Current Opinion in Neurobiology 43 (April): 110–18. 10.1016/j.conb.2017.01.005.

Hammarlund, Marc, Oliver Hobert, David M. Miller, and Nenad Sestan. 2018. “The CeNGEN Project: The Complete Gene Expression Map of an Entire Nervous System.” Neuron 99 (3): 430–33. 10.1016/j.neuron.2018.07.042.

How, Javier J., Saket Navlakha, and Sreekanth H. Chalasani. 2021. “Neural Network Features Distinguish Chemosensory Stimuli in Caenorhabditis Elegans.” PLoS Computational Biology 17 (11): e1009591. 10.1371/journal.pcbi.1009591.

Itskovits, Eyal, Rotem Ruach, Alexander Kazakov, and Alon Zaslaver. 2018. “Concerted Pulsatile and Graded Neural Dynamics Enables Efficient Chemotaxis in C. Elegans.” Nature Communications 9 (1): 2866. 10.1038/s41467-018-05151-2.

Iwanir, Shachar, Rotem Ruach, Eyal Itskovits, Christian O. Pritz, Eduard Bokman, and Alon Zaslaver. 2019. “Irrational Behavior in C. Elegans Arises from Asymmetric Modulatory Effects within Single Sensory Neurons.” Nature Communications 10 (1): 3202. 10.1038/s41467-019-11163-3.

J. Fan, L. Ding, Y. Chen, M. Udell. 2019. “Factor Group-Sparse Regularization for Efficient Low-Rank Matrix Recovery.” Advances in Neural Information Processing Systems 32.

Jin, Eugene Jennifer, Seungmee Park, Xiaohui Lyu, and Yishi Jin. 2020. “Gap Junctions: Historical Discoveries and New Findings in the Caenorhabditis Elegans Nervous System.” Biology Open 9 (8): bio053983. 10.1242/bio.053983.

Kaplan, J M, and H R Horvitz. 1993. “A Dual Mechanosensory and Chemosensory Neuron in Caenorhabditis Elegans.” Proceedings of the National Academy of Sciences of the United States of America 90 (6): 2227–31.

Kato, Saul, Yifan Xu, Christine E. Cho, L. F. Abbott, and Cornelia I. Bargmann. 2014. “Temporal Responses of C. Elegans Chemosensory Neurons Are Preserved in Behavioral Dynamics.” Neuron 81 (3): 616–28. 10.1016/j.neuron.2013.11.020.

Krzyzanowski, Michelle C., Chantal Brueggemann, Meredith J. Ezak, Jordan F. Wood, Kerry L. Michaels, Christopher A. Jackson, Bi-Tzen Juang, et al. 2013. “The C. Elegans cGMP-Dependent Protein Kinase EGL-4 Regulates Nociceptive Behavioral Sensitivity.” PLOS Genetics 9 (7): e1003619. 10.1371/journal.pgen.1003619.

Leinwand, Sarah G., and Sreekanth H. Chalasani. 2013. “Neuropeptide Signaling Remodels Chemosensory Circuit Composition in Caenorhabditis Elegans.” Nature Neuroscience 16 (10): 1461–67. 10.1038/nn.3511.

Leinwand, Sarah G, Claire J Yang, Daphne Bazopoulou, Nikos Chronis, Jagan Srinivasan, and Sreekanth H Chalasani. 2015. “Circuit Mechanisms Encoding Odors and Driving Aging-Associated Behavioral Declines in Caenorhabditis Elegans.” Edited by Oliver Hobert. eLife 4 (September): e10181. 10.7554/eLife.10181.

Lin, Albert, Shanshan Qin, Helena Casademunt, Min Wu, Wesley Hung, Gregory Cain, Nicolas Z Tan, Raymond Valenzuela, Leila Lesanpezeshki, and Vivek Venkatachalam. 2023. “Functional Imaging and Quantification of Multineuronal Olfactory Responses in C. Elegans.” Science Advances 9 (9): eade1249.

Lubba, Carl H., Sarab S. Sethi, Philip Knaute, Simon R. Schultz, Ben D. Fulcher, and Nick S. Jones. 2019. “Catch22: CAnonical Time-Series CHaracteristics.” Data Mining and Knowledge Discovery 33 (6): 1821–52. 10.1007/s10618-019-00647-x.

Metaxakis, Athanasios, Dionysia Petratou, and Nektarios Tavernarakis. 2018. “Multimodal Sensory Processing in Caenorhabditis Elegans.” Open Biology 8 (6): 180049. 10.1098/rsob.180049.

Nguyen, Jeffrey P., Frederick B. Shipley, Ashley N. Linder, George S. Plummer, Mochi Liu, Sagar U. Setru, Joshua W. Shaevitz, and Andrew M. Leifer. 2016. “Whole-Brain Calcium Imaging with Cellular Resolution in Freely Behaving Caenorhabditis Elegans.” Proceedings of the National Academy of Sciences 113 (8): E1074–81. 10.1073/pnas.1507110112.

Prasad, Brinda C, and Randall R Reed. 1999. “Chemosensation: Molecular Mechanisms in Worms and Mammals.” Trends in Genetics 15 (4): 150–53. 10.1016/S0168-9525(99)01695-9.

Pritz, Christian, Eyal Itskovits, Eduard Bokman, Rotem Ruach, Vladimir Gritsenko, Tal Nelken, Mai Menasherof, Aharon Azulay, and Alon Zaslaver. 2023. “Principles for Coding Associative Memories in a Compact Neural Network.” Edited by Yuichi Iino and Timothy E Behrens. eLife 12 (May): e74434. 10.7554/eLife.74434.

Randi, Francesco, and Andrew M Leifer. 2020. “Measuring and Modeling Whole-Brain Neural Dynamics in Caenorhabditis Elegans.” *Current Opinion in Neurobiology*, Whole-brain interactions between neural circuits, 65 (December): 167–75. 10.1016/j.conb.2020.11.001.

Ruach, Rotem, Shai Yellinek, Eyal Itskovits, Noa Deshe, Yifat Eliezer, Eduard Bokman, and Alon Zaslaver. 2022. “A Negative Feedback Loop in the GPCR Pathway Underlies Efficient Coding of External Stimuli.” Molecular Systems Biology 18 (9): e10514. 10.15252/msb.202110514.

Schrödel, Tina, Robert Prevedel, Karin Aumayr, Manuel Zimmer, and Alipasha Vaziri. 2013. “Brain-Wide 3D Imaging of Neuronal Activity in Caenorhabditis Elegans with Sculpted Light.” Nature Methods 10 (10): 1013–20. 10.1038/nmeth.2637.

Sengupta, Piali. 2007. “Generation and Modulation of Chemosensory Behaviors in C. Elegans.” Pflügers Archiv - European Journal of Physiology 454 (5): 721–34. 10.1007/s00424-006-0196-9.

Simonsen, Karina T., Donald G. Moerman, and Christian C. Naus. 2014. “Gap Junctions in C. Elegans.” Frontiers in Physiology 5: 40. 10.3389/fphys.2014.00040.

Small, Dana M., and Barry G. Green. 2012. “A Proposed Model of a Flavor Modality.” In The Neural Bases of Multisensory Processes, edited by Micah M. Murray and Mark T. Wallace. Frontiers in Neuroscience. Boca Raton (FL): CRC Press/Taylor & Francis. http://www.ncbi.nlm.nih.gov/books/NBK92876/.

Suzuki, Hiroshi, Tod R. Thiele, Serge Faumont, Marina Ezcurra, Shawn R. Lockery, and William R. Schafer. 2008. “Functional Asymmetry in Caenorhabditis Elegans Taste Neurons and Its Computational Role in Chemotaxis.” Nature 454 (7200): 114–17. 10.1038/nature06927.

Taylor, Seth R., Gabriel Santpere, Alexis Weinreb, Alec Barrett, Molly B. Reilly, Chuan Xu, Erdem Varol, et al. 2021. “Molecular Topography of an Entire Nervous System.” Cell 184 (16): 4329–4347.e23. 10.1016/j.cell.2021.06.023.

Toyoshima, Yu, Terumasa Tokunaga, Osamu Hirose, Manami Kanamori, Takayuki Teramoto, Moon Sun Jang, Sayuri Kuge, Takeshi Ishihara, Ryo Yoshida, and Yuichi Iino. 2016. “Accurate Automatic Detection of Densely Distributed Cell Nuclei in 3D Space.” PLOS Computational Biology 12 (6): e1004970. 10.1371/journal.pcbi.1004970.

Troemel, E. R., A. Sagasti, and C. I. Bargmann. 1999. “Lateral Signaling Mediated by Axon Contact and Calcium Entry Regulates Asymmetric Odorant Receptor Expression in C. Elegans.” Cell 99 (4): 387–98.

Varshney, Lav R., Beth L. Chen, Eric Paniagua, David H. Hall, and Dmitri B. Chklovskii. 2011. “Structural Properties of the Caenorhabditis Elegans Neuronal Network.” PLOS Computational Biology 7 (2): e1001066. 10.1371/journal.pcbi.1001066.

Wes, P. D., and C. I. Bargmann. 2001. “C. Elegans Odour Discrimination Requires Asymmetric Diversity in Olfactory Neurons.” Nature 410 (6829): 698–701. 10.1038/35070581.

White, J. G., E. Southgate, J. N. Thomson, and S. Brenner. 1986. “The Structure of the Nervous System of the Nematode Caenorhabditis Elegans.” *Philosophical Transactions of the Royal Society of London. Series B*, Biological Sciences 314 (1165): 1–340.

Wilson, Christopher D., Gabriela O. Serrano, Alexei A. Koulakov, and Dmitry Rinberg. 2017. “A Primacy Code for Odor Identity.” Nature Communications 8 (1): 1477. 10.1038/s41467-017-01432-4.

Witvliet, Daniel, Ben Mulcahy, James K. Mitchell, Yaron Meirovitch, Daniel R. Berger, Yuelong Wu, Yufang Liu, et al. 2021. “Connectomes across Development Reveal Principles of Brain Maturation.” Nature 596 (7871): 257–61. 10.1038/s41586-021-03778-8.

Yemini, Eviatar, Albert Lin, Amin Nejatbakhsh, Erdem Varol, Ruoxi Sun, Gonzalo E. Mena, Aravinthan D. T. Samuel, Liam Paninski, Vivek Venkatachalam, and Oliver Hobert. 2021. “NeuroPAL: A Multicolor Atlas for Whole-Brain Neuronal Identification in C. Elegans.” Cell 184 (1): 272–288.e11. 10.1016/j.cell.2020.12.012.

Yu, Yanxun V., Weikang Xue, and Yuanhua Chen. 2022. “Multisensory Integration in Caenorhabditis Elegans in Comparison to Mammals.” Brain Sciences 12 (10): 1368. 10.3390/brainsci12101368.

Zaslaver, Alon, Idan Liani, Oshrat Shtangel, Shira Ginzburg, Lisa Yee, and Paul W. Sternberg. 2015. “Hierarchical Sparse Coding in the Sensory System of Caenorhabditis Elegans.” Proceedings of the National Academy of Sciences of the United States of America 112 (4): 1185–89. 10.1073/pnas.1423656112.

Zufall, Frank, and Steven D. Munger. 2016. Chemosensory Transduction: The Detection of Odors, Tastes, and Other Chemostimuli. Academic Press.

